# MInt-HDX: Leveraging Hydrogen-Deuterium Exchange Mass Spectrometry and Machine-Learning to Improve Protein-Ligand Docking

**DOI:** 10.64898/2026.07.14.738285

**Authors:** Vincent M. Lowe, Ally K. Smith, Rinky Parakra, Eucolona Toci, Caren Freel Meyers, Daniel Deredge

## Abstract

Understanding protein structural dynamics is central to elucidating biological function and guiding therapeutic discovery. Hydrogen-deuterium exchange mass spectrometry (HDX-MS) typically offers peptide-level, and sometimes residue-level, time-dependent insights into protein structure, conformational dynamics and/or ligand binding. Yet, translating HDX-MS data into atomic-resolution insights and deriving mechanistic understanding remains a key challenge. Integrative strategies which utilize HDX-MS to inform computational modeling or simulations, traditionally leverage HDX-MS data with physics-based approaches through the calculation of protection factors models. Here, we developed MInt-HDX, a hybrid physics-based, machine-learning framework trained on differential HDX-MS signatures across 11 protein-ligand systems or 1032 individual peptides, using eXtreme Gradient Boosting (XGBoost) to guide small-molecule ligand docking and pose selection. By leveraging XGBoost-predicted interacting residues with three-dimensional clustering and convex-hull geometric algorithms, MInt-HDX first generates HDX-guided candidate docking sites in 3D for physics-based molecular docking and then, following docking, employs HDX-MS-informed XGBoost filtering and scoring functions for ligand-pose ranking. MInt-HDX was validated across 3 protein-ligand systems, consistently resulting in Ligand-RMSD within 3 Å of the crystallographic ligand conformation, individual steps of MInt- HDX were optimized and its overall performance was assessed against HDX-MS data quality factors and benchmarked against common physics-based and machine learning based docking approaches. Together, this work highlights how machine learning, informed by HDX-MS and aided by physics-based approaches, can bridge the gap between solution-phase HDX-MS data and structural modeling to accelerate protein-ligand discovery pipelines.

**Graphical Abstract:** 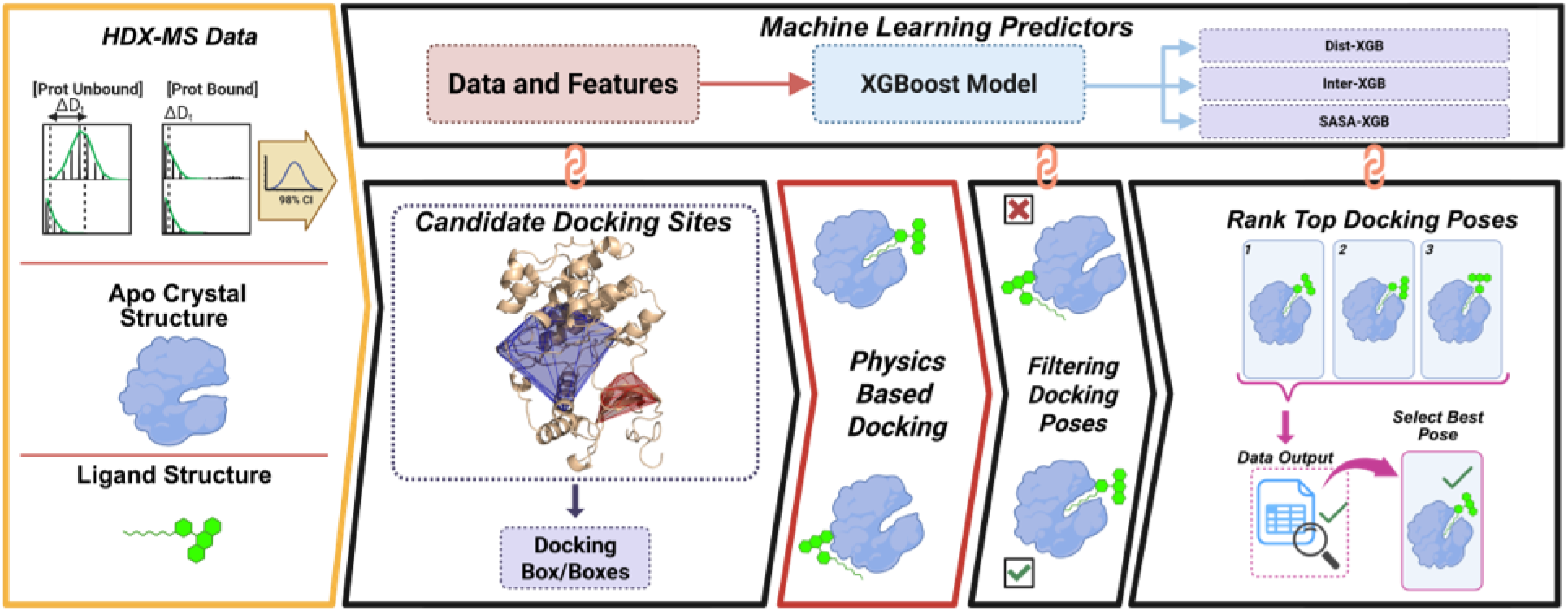

## 1 Introduction

Small-molecule drugs remain a major class of pharmaceutical therapeutics currently in development. In 2025 alone, around 60% of the FDA approved new therapeutics fell into the category of small-molecule drugs.^1^ In addition, many physiological and biochemical processes involve interactions between proteins and small molecules. Therefore, the determination of high-resolution protein-small-molecule ligand complexes is key in understanding both the mechanism of physiological phenomena and in the design of novel small-molecule drugs.^2^ To elucidate the three-dimensional structure of these complexes, a multitude of experimental techniques such as X-Ray crystallography,^3^ cryogenic electron microscopy (Cryo-EM),^4^ or nuclear magnetic resonance (NMR) spectroscopy^5^ have been employed. However, it is not always feasible to obtain an experimentally determined high-resolution structure of the protein-ligand complex,^6–9^ leading to a need for accurate and empirically based computational methodologies.^10^

One such method is simulated docking or molecular docking, in which a small-molecule ligand is placed throughout a search space and the most favorable binding poses are determined through the use of a scoring function.^11^ Numerous molecular docking software tools are available for free or for a fee, such as but not limited to: AutoDock Vina,^12,13^ Rosetta Dock,^14^ Swiss Dock,^15–18^ and Haddock.^19^ However, while many docking tools are available for generating high-resolution structures of protein-ligand complexes, the predicted ligand-binding sites in top-ranking poses often conflict with experimental observations.^20^ In fact, multiple studies have shown the success rate of such tools averaging around 40%, in which success was set as the top pose being within 3 Å of the experimentally determined position of the ligand.^21–23^

With the advent and continued improvement of machine learning-based (ML-based) methodologies, the landscape of structure prediction and determination has been revolutionized. However, even with the vast improvements from each iterative release of software tools such as AlphaFold^24^ and RosettaFold^25^, protein-ligand complexes are still challenging to consistently predict through structure-based prediction models, especially when the protein of interest diverges from the proteins within the dataset.^26–28^

Alternative approaches involve using lower resolution experimental methodologies integrated with computational, physics-based approaches which can provide insight into structural or biological factors of proteins, allowing researchers to better understand how these proteins act in vivo/in vitro. One such experimental method is hydrogen–deuterium exchange coupled with mass spectrometry (HDX-MS).^29^ HDX-MS monitors the exchange of backbone amide hydrogen atoms with deuterium, a stable hydrogen isotope, upon exposure to a deuterated solvent. Using mass spectrometry, this exchange can be monitored at peptide-level resolution, allowing for analysis of solution dynamics as a function of labeling time, typically seconds to hours.^30^ HDX-MS provides information-rich data regarding the protein’s structure and dynamics, resolved in both space and time. Previous work has illustrated the promise of HDX-MS data’s integration with computational approaches such as with quantitative correlation,^31,32^ ensemble reweighting,^33,34^ and even protein-ligand docking.^35,36^ Furthermore, the availability of HDX-MS data has only grown in the past years with databases such as the PRIDE (Proteomics Identifications Database)^37^ and MassIVE (Mass Spectrometry Interactive Virtual Environment)^38^ databases. The increasing availability of HDX-MS data has led to a multitude of experimentally trained computational methods including but not limited to AI-HDX,^39^ HDXRank,^40^ HDXmodeller.^41^ Each of these advancements have shown the value of integrating experimental HDX-MS data with computational methodologies and algorithms to further delve into the structure, features, and factors that influence proteins in solution.

Here, we have developed and benchmarked a generalizable machine-learning approach known as Modeling with Integrative HDX-MS (MInt-HDX) that utilizes HDX-MS experimental data to guide and automate protein-small-molecule ligand docking, and to filter and score the docking poses generated. Built on the foundation of a supervised eXtreme Gradient Boosting (XGboost) algorithm,^42,43^ MInt-HDX was trained and validated on a total of 14 datasets curated from both our in-house and the PRIDE databases,^37^ with both the bound and unbound crystal structures and the HDX-MS data. After training and validating our workflow, we compared the results of MInt- HDX to the current gold standards in molecular docking,^20^ as well as explored the effects of the quality, redundancy, and coverage of the HDX-MS data on the algorithms predictive and scoring capability. Our results illustrate the power of HDX-MS data integrated with ML in guiding, automating, and ranking the docking poses for protein-small-molecule docking, even in the presence of a limited dataset.

## 2 Methodology

### 2.1 HDX-MS Data Curation

While an HDX-MS dataset can be highly complex and information rich, one major hurdle to the prediction of protein-ligand docking poses is the limited amount of well-curated, publicly-available datasets. To help address this issue, we searched and curated through our in-house and the PRIDE^37^ databases for datasets that satisfied the following criteria: the presence of experimentally determined high-resolution structures of the protein in both the ligand-bound and unbound states, a known structure of the ligand, and the HDX-MS data with deuterium uptake and uptake standard deviation information for both the bound and unbound states. With these inclusion criteria, we were able to collect 14 datasets from both the PRIDE database and our in-house datasets. From the HDX-MS data, we calculated the deuteration difference (ΔD) of the bound minus the unbound state’s absolute deuterium uptake for each peptide at every exposure time. To reduce the noise within the datasets and ensure the model was trained on significant HDX-MS values, we applied a 98% two-sample pooled T confidence interval calculated as shown in Houde et al.,^44,45^ setting any value that fell within the confidence interval, and was therefore deemed to be non-significant, to zero. The data were split into 11 training and 3 validation datasets (Supplemental Tables S1 and S2) while ensuring good distribution of protein type and size across training and validation sets. The 11 training data sets amounted to a total of 1032 individual peptides with differential HDX-MS data.

### 2.2 XGBoost Model Training

We trained three separate XGBoost models to predict three separate structural features that provide information on protein-ligand binding: Interaction (Inter-XGB), Distance (Dist-XGB), and solvent accessible surface area (SASA-XGB). Prior to training the three models, the target features for each model were calculated from the available crystal structures and transformed into binary classification values. For the Inter-XGB model, a program known as the Protein-Ligand Interaction Profiler (PLIP) was used, with default parameters, to obtain the residues in the bound crystal structure that were interacting with the ligand.^46^ Due to the peptide-level resolution of HDX-MS data, the residue-level interactions were extrapolated to the peptide level by assigning an entire peptide segment to be interacting if a residue within a peptide segment was determined to be interacting with the ligand. For the Dist-XGB model, a target feature providing distance insights into the protein-small-molecule interaction was used, classifying any residue as interacting if any atom of a given residue is within a certain distance of any atom of the ligand. The distances were obtained using Pymol’s python scripting package, and the “within” function of said package.^47^ The interaction distance was optimized to a default value of 5 Å of any atom on the ligand. Again, this information was extrapolated from residue level to peptide level. For the SASA-XGB model, residue-level decreases in SASA upon ligand binding were calculated. For the purposes of a binary classification, we optimized a threshold value of ≥ 50% decrease in their SASA, between the bound and unbound states, as interacting. To obtain SASA, the Shrake Rupley method was used within the Biopython python package.^48,49^ SASA calculations of the unbound state were performed with the empirically determined unbound structure as well as the bound structure minus the ligand. However, to ensure this assumption would be valid in comparison to using the crystal structure of the unbound, we calculated the RMSD between the unbound and bound structures within the datasets. The results of these calculations can be found in Supplemental Table S3. Here again, the residue level results were extrapolated to the peptide level.

The training of the three XGBoost models were performed using the Scikit-Learn python package.^50^ For that purpose, the ΔD data of the training set, pre-processed through the 98% statistical confidence interval, was used as the input data for the XGBoost package. The hyperparameters for each model were tuned using the python package Optuna^51^ (Supplemental Figure S1 and Supplemental Tables S4, S5 and S6). Once trained and optimized, these models were then leveraged at various stages of MInt-HDX, a hybrid physics-based/ML-based workflow which includes 1/ defining candidate docking sites, 2/ performing physics-based molecular docking, 3/ filtering out poses docked at distal sites, and 4/ scoring and ranking docking poses. The trained XGBoost models were leveraged in steps 1, 3 and 4.

### 2.3 Defining Candidate Ligand Docking Site using HDX-trained XGBoost Models

To define candidate docking sites, the Inter-XGB and Dist-XGB models were used to extract all the peptide segments predicted by both models to be interacting with the ligand. The coordinates of the alpha carbons of these peptide segments were then mapped in 3-dimensional space. Density Based Spatial Cluster of Applications with Noise (DBSCAN) which clusters points using a density-based notion in 3-dimensional space, was used to cluster these alpha carbons coordinate points.^50,52^ From there, convex hull theory, which determines the smallest convex polygon that contains any given set of points in n-dimensions of space, was used to construct a polygonal shape from the resulting clusters of alpha carbon points in 3D space.^53^ These polygonal shapes represent a hotspot of alpha carbon atoms from peptides predicted to be interacting and were used as a candidate docking site for downstream docking. The size and number of these candidate docking sites is dependent upon a confidence threshold value which filters out peptide segments based on the probability of the peptide interacting, as determined by both the Inter- XGB and Dist-XGB. A default confidence threshold of 0.7 or 70% probability was set, as determined through screening the confidence threshold values.

### 2.4 Molecular Docking to the Candidate Docking Sites

Using an internal code, the smallest box was drawn around each individual candidate docking site to generate a docking box required by Autodock Vina.^12^ To maximize the probability of producing an accurate docking pose, the docking box’s size was checked to determine if all dimensions of the box were ≥ 2.9 times the radius of gyration of the ligand, as defined by Feinstein and Brylinski.^54^ The dimensions of the box were adjusted accordingly, as needed. Autodock Vina was used due to its ability to integrate within a python code pipeline, allowing an automated workflow to be developed for pre-docking preparation and post-docking analysis. Autodock Vina was used with default parameters except for a change to the exhaustiveness^55^ and n_poses, which were set to 50 and 20, respectively. This workflow was performed for each candidate docking site and poses were generated for each individual site. Poses which correspond to the most likely site were then determined through a filtering function as described in section 2.5.

### 2.5 MInt-HDX Filtering Function

Dist-XGB was used to incorporate a filtering function following the generation of docking poses and prior to the scoring step. The filtering function serves the purpose of filtering poses that deviate significantly from the crystallographic ligand pose. For instance, such poses can result from distal candidate docking site that can arise from long range or allosteric HDX-MS protection that can occasionally be observed upon ligand binding. The equation for filtering is shown in equation 1:

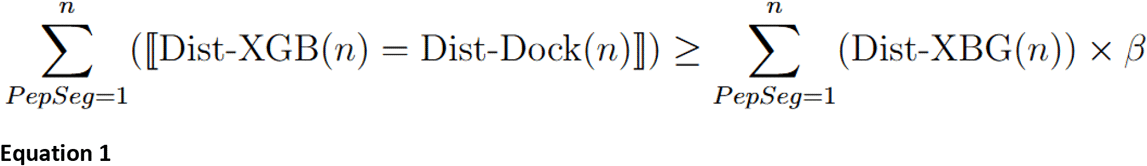

Where PepSeg is the peptide segments, Dist-XGB(n) is a Dist-XGB predicted binary value of interaction (0 or 1) for a given peptide segment n, and Dist-Dock is the observed binary value of interaction (0 or 1) for peptide segment n in a given docking pose Here, interaction is defined as any atom of the peptide segment being within 5 Å of the docked ligand, and [[…]] is an Iverson bracket, which equals 1 when the stated condition is true and 0 when it is false. The equation filters out docking poses where the agreement between Dist-XGB and Dist-Dock falls below a certain fraction (β coefficient) of all Dist-XGB predicted interacting peptides. The β coefficient was systematically benchmarked in the validation dataset to optimize filtering out poses that are more than 10 Å away from the true binding site.

### 2.6 MInt-HDX Scoring Function

The poses retained by the filtering function were then scored and ranked based on a scoring function Score_Avg_ (Equation 2) that combines scoring functions from all three individual XGBoost models, Score_Int_, Score_Dist_ and Score_ΔSASA_. Individual scoring functions were calculated as in Equation 3A, 3B and 3C, where PepSeg are the peptide segments, Inter-XBG, Dist-XGB and SASA- XGB are the respective predicted binary value of interaction (0 or 1) for a given peptide segment n; Inter-Dock, Dist-Dock and SASA-Dock are the observed binary values of interaction extracted from the docking poses using the respective established definition of interaction, and [[…]] is an Iverson bracket, which equals 1 when the stated condition is true and 0 when it is false. Only peptide segments with XGBoost prediction values with probabilities ≥ 0.7 were considered when scoring, to ensure higher confidence in the selected poses.

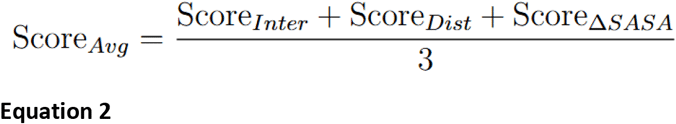

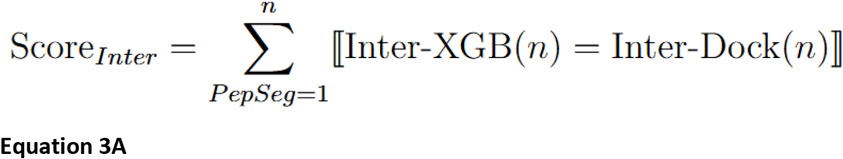

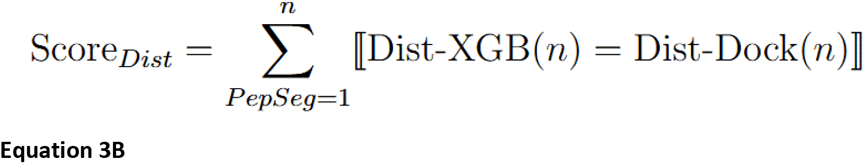

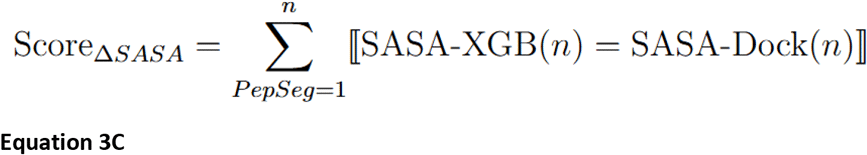

Docking poses with the higher Score_avg_ were deemed to be the better models of the protein-ligand complex. When two poses had identical Score_Avg_ values, Autodock Vina’s scoring function was used to determine rank.

### 2.7 Benchmarking Against Alternative Docking Approaches

To compare the performance of MInt-HDX enabled docking vs. commonly used Molecular Docking tools (AutoDock Vina^12,13^ and SwissDock^15–18)^, we performed molecular docking following each docking tool’s documented workflow for the three systems in our validation set in triplicate. For Autodock Vina, we generated the docking box by using AutoGrid4, provided by the AutoDock Suite. AutoGrid4 calculated the grid maps that Vina used during the ligand docking search process.^56^ Additionally, we also compared MInt_HDX to DiffDock, an entirely machine learning based, generative, docking tool. DiffDock runs were performed in triplicate with default settings.^57,58^

The accuracy of docking poses from MInt-HDX, AutoDock Vina, SwissDock and DiffDock were assessed against the crystal structure conformation as follows. For any given docking pose, the protein was structurally aligned, using Pymol’s align function,^47^ to the ligand-bound co-crystal structure using backbone atoms. Then the all-atom RMSD of the ligand in the docking pose to the crystal structure ligand (L-RMSD) was calculated and used as a metric of docking accuracy. For each method, the top one (Top-1) and top three (Top-3) poses were selected to compare L- RMSD and assess the performance and accuracy of MInt-HDX and other docking workflows.

### 2.8 Benchmarking Performance Against HDX-MS Data quality: Average Peptide Coverage Redundancy and Average Peptide Length

To explore the effect of the HDX-MS data quality on the performance of MInt-HDX, we benchmarked two key factors pertaining to the HDX-MS data quality: average peptide coverage redundancy and average peptide length. Redundancy refers to how many unique peptide segments cover a specific residue, with average peptide coverage redundancy referring to the peptide coverage redundancy averaged over all residues. Average peptide length is interpreted more as a reflection of the efficiency of the acid-protease digestion of the protein into peptide segments and as an indicator of the resolution of the HDX-MS data obtained. To explore the effect of average peptide length on MInt-HDX’s performance, we artificially trimmed the HDX-MS peptide segment list of our three validation systems to include only peptide segments with a length ≥ 10 residues or ≤ 9 residues. Similarly, to probe the effects of loss of average peptide coverage redundancy, the peptide segment lists were set to have an artificial average peptide coverage redundancy of ∼1, while keeping the average peptide length and the sequence coverage the same as the original data. Each of these datasets was analyzed in triplicate to explore the variance of the results under these conditions.

### 2.9 Test Case: Evaluating the Effect of Missing HDX-MS Sequence Coverage Near the Binding Site with *Dr*DXPS

*Deinococcus radiodurans* 1-deoxy-D-xylulose 5-phosphate synthase (*Dr*DXPS) was overexpressed in *E. coli* BL21 (DE3) cells harboring dxs-pET28b(+)^59^ and purified as previously reported^60^ using an AKTA-GO fast protein liquid-chromatography (FPLC) system. HDX-MS data for *Dr*DXPS was acquired as follows. HDX-MS data of *Dr*DXPS with and without the methyl acetylphosphonate (MAP) ligand was obtained as follows. *Dr*DXPS (12 µM) was exchanged into buffer containing 50 mM HEPES pH 8, 0.2 mM ThDP, 50 mM NaCl, 1 mM MgCl_2_. DXPS was incubated alone or with 1 mM MAP for 20 min at RT before HDX reactions. The experiments were initiated by mixing 2 µL of the DXPS sample with 38 µL of ^2^H_2_O buffer (exchange buffer prepared in 99.9% ^2^H_2_O). The HDX reaction mixtures were incubated for 10 s, 20 s, 1 min, 5 min, 10 min, 1 h, and 2 h and then quenched by rapidly mixing with 40 µL of ice-cold quench buffer (100 mM glycine, 8 M Urea, 20 mM TCEP, pH 2.5). The samples were then immediately flash frozen and stored at -80 ℃. Sequencing controls were prepared similarly but in H_2_O buffer, fully deuterated controls were prepared by denaturing *Dr*DXPS in 7M guanidine, then incubating in ^2^H_2_O exchange buffer for 2 h prior to quenching as described above. MInt-HDX was performed with the resulting data after removing regions displaying EX1 kinetics. Default values were initially used and subsequently, the confidence threshold value was modulated.

### 2.10 Test Case: Evaluating the Ability of MInt-HDX to Identify Favored Binding Conformations of Imatinib with c-src Kinase

To test the ability of MInt-HDX to distinguish between distinct binding conformations of the same ligand, we used c-src kinase binding to Imatinib. Three starting structures of c-src were used for docking. Using the crystal structure of chicken c-src bound to Imatinib in the DFG-out form (PDB:2OIQ), a pseudo-apo structure was generated by removing the ligand from the binding site in chain A. From the same crystal structure, chain B was assessed for DFG conformation and used. Electron density for the ligand is not observed in chain B. Finally, using another crystal structure of the kinase domain of the human c-src bound to MPZ, an Imatinib analog, in DFG-in conformation (PDB: 1Y57), we created another pseudo-apo form by removing the ligand from the binding site. 2OIQ and 1Y57 have a sequence similarity of 99%, with the only difference in the kinase domains being a single residue shift between chain A of 2OIQ and 1Y57. Of note, MInt- HDX was performed with a confidence threshold value of 0.5 due to the absence of candidate docking sites at a value of 0.7.

## 3 Results and Discussion

### 3.1 Assessment and Optimization of the Performance of Individual Steps of MInt-HDX

Prior to assessing the overall performance of our workflow, we independently benchmarked and optimized each step of our workflow that leveraged our trained XGBoost models. The results of each step are shown in Figure 1.

**Figure 1:**
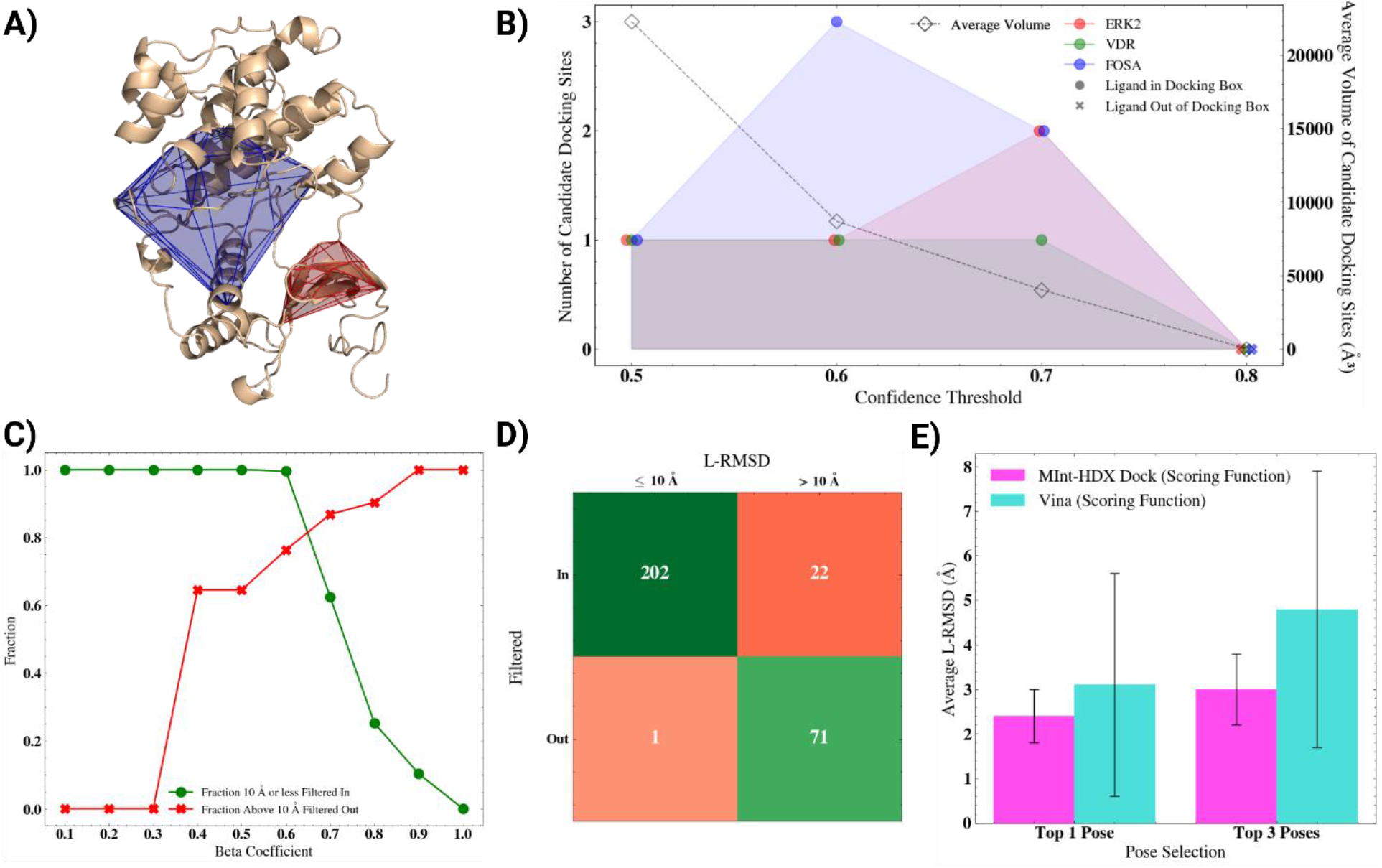
MInt-HDX workflow evaluation and optimization. **A)** The DBSCAN and convex hull algorithms were used to identify and map the candidate docking sites of the ligand to the protein. Shown are two separate candidate docking sites (in blue and red) mapped to the ERK2 protein shown in cartoon representation. **B)** For each of the validation set proteins, the number of candidate docking sites (y-axis 1) and the average volume of the candidate docking sites (y- axis 2) were plotted against the confidence threshold value. The presence of crystallographic ligand binding site within the resulting docking box generated from those sites was mapped based on the marker type. **C)** The optimal Beta Coefficient for filtering. The fraction of poses with L-RMSD ≤ 10 Å and filtered in (Green) or > 10 Å and filtered out (Red) plotted vs the Beta Coefficient. This was performed on all 296 poses generated for the validation of set proteins. **D)** The confusion matrix showing the L-RMSD vs the result of the Filtering for β = 0.6. **E)** Comparison of MInt-HDX and AutoDock Vina’s scoring function applied to the filtered poses. The average L-RMSD and standard deviations of Top-1 and Top-3 poses are plotted in a bar graph according to their respective scoring functions.

#### 3.1.1 Optimized Confidence Threshold for HDX Guided Generation of Candidate Docking Sites

The first step involves generating candidate ligand docking sites in 3D space by leveraging peptides predicted to be interacting by Inter-XGB and Dist-XGB and using a combination of DBSCAN clustering and convex hull theory. To assess the effect of confidence threshold on the ability of Inter-XGB and Dist-XGB to optimally predict interacting peptides for this purpose, the number of candidate docking sites, the average volume of the resulting candidate docking sites and the presence of the crystallographic ligand within the docking boxes was plotted against the confidence threshold as shown in Figure 1B and Supplemental Table S7. Across all proteins in the validation dataset, at confidence thresholds above 0.7, there were not enough points to successfully generate a candidate docking site, which resulted in the unsuccessful completion of the workflow. This reflects the fact that Inter-XGB and Dist-XGB predictions resulted in few or no peptides above 0.75 probability. As the confidence threshold was lowered to 0.7 and below, the number of distinct candidate docking sites generated increased with a maximum between 0.6 and 0.7. As the confidence threshold was decreased further towards 0.5, the candidate sites agglomerated into a single, large site. As a consequence, the average volume of the candidate docking site increased gradually as the confidence threshold decreased from 0.7 to 0.5. While such large candidate docking sites could maximize the likelihood of containing the ligand binding site, they are also likely to include potential alternative sites for AutoDock Vina to dock in the subsequent step. Further inspection revealed that, irrespective of the confidence threshold, the docking boxes resulting from the candidate docking sites always contained the ligand as in the crystallographic pose when aligned with the co-crystallized bound structure. Consequently, a default confidence threshold of 0.7, which maximizes the number of docking boxes but minimizes the docking box volumes, was chosen and used for the assessment of our workflow across the validation set, unless otherwise stated.

#### 3.1.2 Optimized β Coefficient Value for MInt-HDX’s Filtering Function

To determine the optimal β coefficient value, we applied the filtering function to all docking poses generated for the three validation set proteins at a confidence threshold of 0.7. At that confidence threshold, five candidate docking sites were generated across the three proteins. Docking was performed in triplicate for each of the resulting docking box, generating 60 docking poses per candidate docking site as shown in Figure 1B. A total of 296 docking poses were extracted due to variability in output from AutoDock Vina. Across these poses, the β coefficient was varied from 0.1 to 1, as shown in Figure 1C and Supplemental Table S8, and the ability of the filtering function (Equation 1) to accurately filter ligand poses with an L-RMSD greater than 10 Å was assessed. A value of 10 Å was chosen as an arbitrary value reflecting the ability of Equation 1 to effectively filter out docking poses at docking sites distal from the true binding sites, and also to filter out poses of ligands in the correct sites yet in a largely incorrect orientation. The latter is particularly relevant for ligands with significant symmetry. Probing across β coefficient space, a value of 0.6 was observed to be most efficient at filtering out docking poses. Indeed, shown in Figure 1D are the results of the filtering at β = 0.6, with poses classified as having an L-RMSD ≤ 10 Å, or > 10 Å. Our results showed that the filtering function successfully filtered in 99.5% of poses with an L-RMSD of 10 Å or less and filtered out 76.3% of the poses with L-RMSD greater than 10 Å. A default value of β = 0.6 was subsequently used. Of note, the filtering function equation can bias for regions of protection with the highest HDX-MS peptide coverage redundancy and works best under the assumption that the true binding site is populated with equivalent HDX-MS peptide coverage redundancy as alternative sites showing protection from deuterium uptake.

#### 3.1.3 Comparison of MInt-HDX vs AutoDock Vina’s Scoring Function

To assess the performance of MInt-HDX’s scoring function, we scored and ranked the docking poses that were selected by our filtering function (224 poses at β coefficient = 0.6) and compared the accuracy and precision of our scoring function to that of AutoDock Vina using the L-RMSD to the co-crystallized ligand pose. Shown in Figure 1E are the L-RMSD values of the Top-1 scored and Top-3 scored poses resulting from MInt-HDX and Vina’s scoring functions. The L-RMSD values were averaged across the independent triplicate acquisitions of each of the three proteins in the validation set.

For the same set of docking poses, the MInt-HDX scoring function outperformed the Vina scoring functions in terms of accuracy with L-RMSD values of 2.4 Å vs 3.1 Å for the Top-1 scored poses. In addition, the Top-3 L-RMSD reflected the improved accuracy of the MInt-HDX scoring function with L-RMSD values of 3.0 Å vs 4.8 Å (Figure 1E and Supplemental Table S9). Just as importantly, the Mint-HDX scoring function displayed significantly greater precision with a standard deviation of 0.8 Å for MInt-HDX as opposed to 3.1 Å for Vina (Figure 1E and Supplemental Table S9), increasing the confidence in the performance of the MInt-HDX scoring function; this likely reflects the fact that the MInt-HDX scoring function takes advantage of structural measurement in the form of HDX-MS data-trained XGBoost models, whereas the AutoDock Vina scoring function does not leverage any additional data.

### 3.2 Performance of Overall Workflow

Overall, the MInt-HDX workflow resulted in accurate and precise docking poses. Indeed, it produced an average Top-1 pose L-RMSD of 2.4 Å and an average Top-3 L-RMSD pose of 3.0 Å relative to the experimentally determined co-crystallized poses across the three protein-ligand pairs in the validation set (Table 1, Supplemental Table S10). In addition, MInt-HDX demonstrated strong confidence and precision, with standard deviations of 0.6 Å and 0.8 Å for the Top-1 and Top-3 poses respectively (Table 1, Supplemental Table S10). In small-molecule docking, L-RMSD to the crystallographic pose remains the gold-standard metric for benchmarking pose prediction accuracy, with L-RMSD thresholds of ≤2 Å generally considered successful or as a near-native pose, and L-RMSD between 2-3 Å often regarded as acceptable for medicinal chemistry applications, structural interpretation, and lead optimization.^22,61,62^ Accordingly, the MInt-HDX average Top-1 and Top-3 poses fell within the threshold commonly considered structurally and pharmacologically meaningful. Having established that MInt-HDX reached consensus-acceptable standard for small-molecule docking, we next benchmarked it against other established physics-based and ML-based docking approaches which do not leverage HDX-MS data, before comparing it to an approach which leverages a mixture of molecular dynamics simulations, ensemble reweighting, and HDX-MS data.^35^

**Table 1:** MInt-HDX vs other physics-based or ML-based docking approaches. All docking was performed on three protein systems of the validation set, for 3 replicates each. Shown are the L-RMSD of Top-1 pose and Top-3 poses, averaged across all 9 runs with standard deviations. Each method was performed following the documented workflow, as provided by the developers.

|  | Average Top-1 |  | Average Top-3 |  |
| --- | --- | --- | --- | --- |
|  | L-RMSD | SD | L-RMSD | SD |
| HDX Refined Docking |  |  |  |  |
| Mint-HDX | 2.4 | 0.6 | 3.0 | 0.8 |
| Physics Based Docking |  |  |  |  |
| AutoDock Vina | 11.9 | 2.2 | 12.0 | 1.8 |
| SwissDock | 10.4 | 9.4 | 10.8 | 6.6 |
| ML Based Docking |  |  |  |  |
| DiffDock | 2.1 | 0.3 | 2.1 | 0.1 |

#### 3.2.1 Improved Performance vs. Physics-Based Molecular Docking Tools

We evaluated the performance of MInt-HDX in comparison with two commonly-used molecular docking tools, AutoDock Vina and SwissDock. Each docking tool was run for the three protein-ligand pairs in the validation set, with each protein-ligand docking run conducted in triplicate to capture the variability of the tool’s performance. Docking was performed without any *a priori* information regarding the drug binding site. The Top-1 and Top-3 poses were extracted and their accuracies were compared with MInt-HDX using their respective average L-RMSD to the crystallographic ligand conformation. To analyze the variability of the tool’s predictive capabilities, the standard deviation for the L-RMSD was also determined. When compared to MInt-HDX, AutoDock Vina produced poses with L-RMSD that were an average of 9.5 Å and 9.0 Å higher for Top-1 and Top-3 poses, respectively (Table 1, Supplemental Table S10). In addition, MInt-HDX also showed a greater consistency in deriving accurate poses as indicated by the lower standard deviations in L-RMSD of the Top-1/Top-3 poses. To account for any biases resulting from the molecular docking tool used, we also performed the comparison with SwissDock. Similarly, SwissDock produced docking poses with an average 8.0 Å and 7.8 Å increase in L-RMSD over MInt-HDX Dock and significant increase in standard deviation (Table 1, Supplemental Table S10). Since AutoDock Vina is also part of MInt-HDX’s workflow, this indicated that the increase in performance results from the other component of the workflow. Indeed, MInt-HDX leverages additional mid-resolution structural data in the form of HDX-MS data through the use of HDX trained XGBoost models. In turn, these trained models allow more targeted, data-informed docking through the definition of localized candidate docking sites derived from the HDX-MS data, as well as through the HDX-MS data-based filtering step and scoring function that we observed to be more accurate than AutoDock Vina’s scoring function.

Visual inspection of the docking poses confirmed that MInt-HDX outperformed these molecular docking tools (Figure 2, Supplemental Figure S2, S3 and S4). Figure 2 shows the Top-3 poses from the replicate with the lowest average L-RMSD obtained from either MInt-HDX, AutoDock Vina, SwissDock or DiffDock for each protein-ligand pair in the validation set. Inspection of the docking poses revealed a clear difference between the commonly used search-based tools and MInt-HDX. MInt-HDX was consistently able to accurately identify the binding pocket and produce docking poses in a correct orientation within the binding pocket. On the other hand, the search-based tools frequently docked at least one ligand in the wrong binding site (AutoDock Vina-FosA, SwissDock-Erk2 and FosA) or docked the ligands in the correct site but with markedly incorrect orientations (AutoDock Vina-Erk2 or SwissDock-FosA). It is worth noting that FosA is a symmetrical homodimer with two ligand binding sites. MInt-HDX consistently favored one of the binding sites, whereas AutoDock Vina did not return a docking pose within either of the two sites, and SwissDock returned a docking pose in each of the sites, albeit with limited conformational accuracy. Closer inspection revealed that the MInt-HDX bias to one of the binding sites is likely due to slight differences in local conformation between the binding pockets in the unbound FosA structure. Indeed, in the binding pocket that was favored by MInt-HDX, chain P, the sidechain and backbone conformation mimicked that of the ANY1 co-crystallized pocket conformation. However, in the other binding pocket, chain G, the sidechains of residues E98 and Y131 were turned inwards, altering the local conformation of the binding pocket, which would result in steric hindrance with ANY1 in the co-crystallized conformation. The structural alignment of the disfavored site to the co-crystallized structure revealed an RMSD of 4.8 Å and 1.6 Å for E98 and Y131 respectively. Interestingly, the MInt-HDX workflow generated candidate docking sites for both pockets, docked within each, and filtered in poses from both chains. However, the scoring function favored the chain in which the sidechain conformation is akin to the bound form. This demonstrated the sensitivity of MInt-HDX in successfully discriminating different docking poses resulting from minor differences in conformation of the binding pocket. However, it also points to one limitation of MInt-HDX: it is currently limited to rigid docking. Indeed, with the inclusion of a rigid docking protocol (AutoDock Vina) in MInt-HDX, the need for conformational re- orientation, like that seen in FosA, induced-fit or co-folding can be a limiting factor to accurate docking.

**Figure 2:**
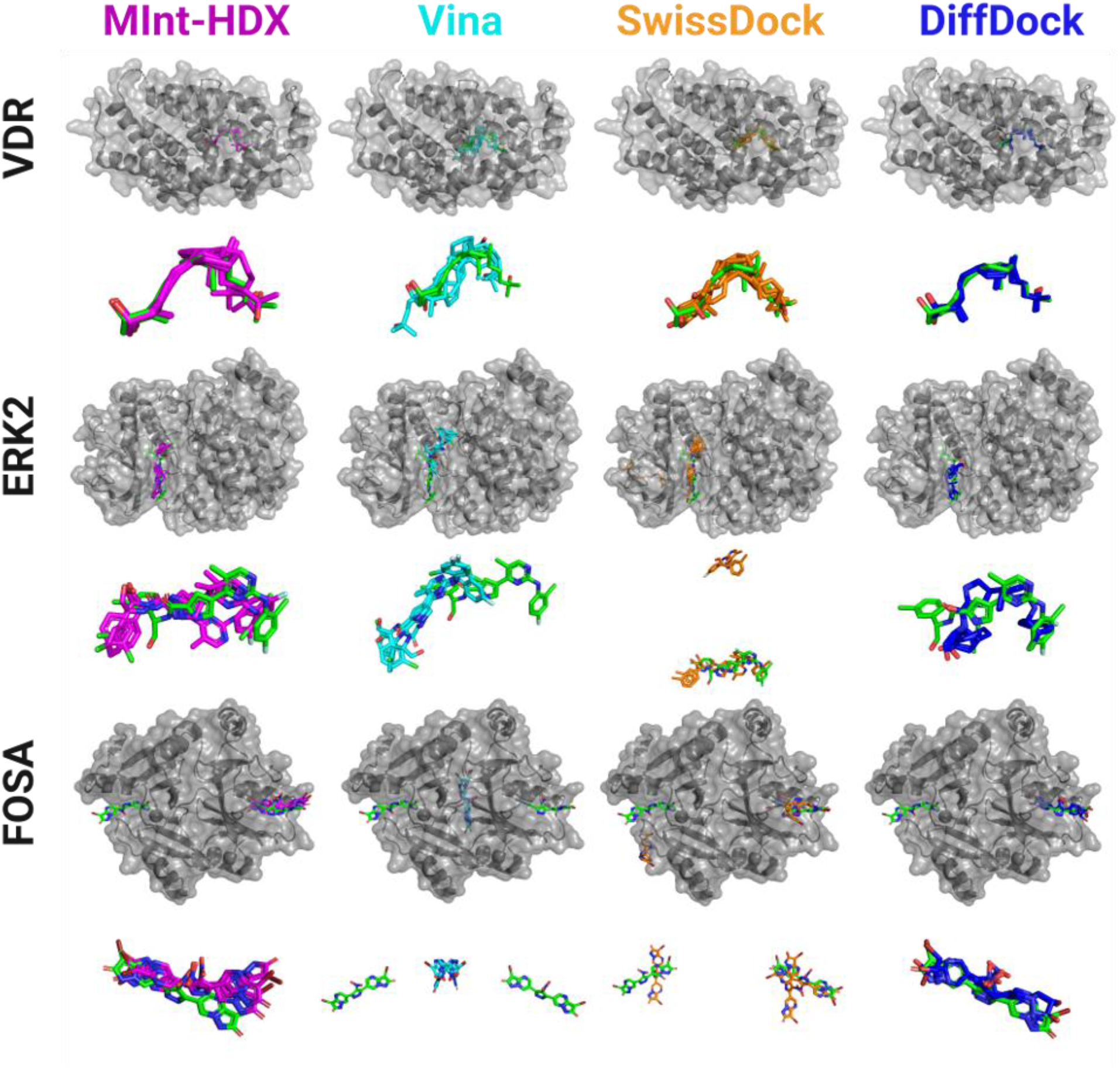
Alignment of the top replicate for each docking approach, across the 3 validation set proteins. The docking poses from MInt-HDX (Magenta), AutoDock Vina (Cyan), SwissDock (Orange), and DiffDock (Blue) were aligned to the ligand from the bound crystal structure (Green). Alignment is performed based on protein backbone atoms. Shown are the Top-3 docking poses from the best replicate of each docking approach by L-RMSD, per protein. The top shows the zoomed out, full protein view with the ligands in sticks and protein in surface representation. The bottom images illustrate the ligand conformation zoomed in, without the protein. The PDB complexes, in order of rows (top to bottom) are 1DB1, 6OPK, and 5WEW. For the bottom left and bottom right zoomed in view of the ligands, the second ligand was left out of the image due to the predictions only being to one part of the homodimer.

#### 3.2.2 Comparison to a Machine Learning-Based Docking Tool

We also evaluated the performance of MInt-HDX in comparison to a ML-based small-molecule docking tool. To that end, we performed docking using the diffusion-based docking tool DiffDock^57,58^ on each of the validation set proteins, in triplicate to capture the variance of the tools. The results indicated comparable outcomes, with both yielding docking poses with an L- RMSD between 2-3 Å from the co-crystallized ligand conformation, although DiffDock slightly outperformed MInt-HDX. Indeed, when comparing the average Top-3 poses resulting from MInt- HDX vs. DiffDock, MInt-HDX displayed an average of 0.9 Å greater L-RMSD than DiffDock. When comparing the average Top-1 poses, MInt-HDX resulted in an average of 0.3 Å greater L-RMSD than DiffDock (Table 1, Supplemental Table S10). DiffDock also yields results with slightly higher precision, with standard deviations in the average Top-1 and Top-3 poses of 0.3 Å and 0.1 Å, respectively. This is also confirmed in the visual inspection of the poses (Figure 2, Supplemental Figures S2, S3 and S4) where DiffDock identifies the binding sites accurately and yields poses in the correct orientation. However, detailed inspection of the performance for individual proteins in the validation set (Supplemental Table S10) revealed that DiffDock outperformed MInt-HDX only for VDR (L-RMSD Top-1 of 0.6 Å vs 2.0 Å), with similar performance for FosA (L-RMSD Top-1 of 2.2 Å vs 2.1 Å) and marginally poorer performance for ERK2 (L-RMSD Top-1 of 3.6 Å vs 3.1 Å), an observation that can be explained by the presence of FosA and VDR in DiffDock’s training data.^57,58^ Regardless, the overall comparable performance is noteworthy in light of the large differences in training dataset size. Indeed, MInt-HDX was trained on 11 protein-ligand complexes. In contrast, DiffDock was trained on 17,000 complexes, meaning that MInt-HDX had a total of 0.06% of DiffDock’s training dataset.^57^ This illustrates that HDX-MS data, a mid-resolution structural approach, captures sufficient structural information to inform a hybrid physics-based, ML-based approach to extract ligand binding poses with accuracy on par with state-of-the-art ML-based docking approaches that benefit from vastly larger dataset to train on. This highlights the potential of integrative HDX approaches that aim to leverage HDX-MS data for modeling and simulations through physics-based and/or ML approaches.

#### 3.2.3 Improved Efficiency Compared to HDX Ensemble Reweighting Approach

We previously developed an integrative HDX approach that used physics-based approaches to model protein-small-molecule interactions. Similar to MInt-HDX, the integrative HDX approach required three inputs, the crystal structure of the protein, the structure of the ligand, and the ligand-bound and unbound HDX-MS data. It consisted of a workflow that employs a combination of molecular docking, molecular dynamics simulations, a *post hoc* hydrogen-deuterium exchange ensemble reweighting (HDXer)^33,63^ method, and dimensional reduction and clustering approaches to extract docking poses that were guided by experimental HDX-MS data.^35^ One of the protein systems used in that study, FosA binding to ANY1, was included in the validation set of MInt-HDX, allowing for a direct comparison of performance of the two methods.

The application of the HDXer-based workflow resulted in a docking pose with a L-RMSD of 2.0 Å.^35^ Comparatively, MInt-HDX resulted in an average Top-1 pose for FosA-ANY1 with a L-RMSD of 2.1 Å (Supplemental Table S10). While MInt-HDX and our HDXer based workflow achieved similar accuracy for the FosA-ANY1 interaction, the computational cost and time required to execute both methods are substantially different. The reweighting workflow, which includes docking, molecular dynamics simulations, ensemble reweighting, and dimensional reduction and clustering, typically takes several days to weeks to complete. In contrast, MInt-HDX operates on a timescale of minutes to hours with molecular docking through AutoDock Vina as the primary bottleneck. These results highlight critical gains in efficiency and rapid turnaround by MInt-HDX that would allow accelerated performance and scalability, and ultimately enable envisioning the incorporation of integrative HDX approaches in high-throughput small-molecule screening efforts by HDX-MS.

#### 3.2.4 HDX-MS Data Quality can Affect MInt-HDX’s Performance

Typical experimental factors that affect the quality and interpretability of HDX-MS data include the average peptide length and average peptide coverage redundancy. The average peptide length is considered to be indicative of the resolution of the HDX-MS data. Peptide segments which are made up of a higher number of residues and are therefore longer will be lower resolution due to the difficulty in parsing out the effects of individual residues. In contrast, peptide segments which are made up of fewer residues and are therefore shorter, will have a higher resolution, allowing for a better understanding of the individual effects. The peptide coverage redundancy is the number of peptides covering a given residue. The average peptide coverage redundancy is the peptide coverage redundancy averaged over all residues and is considered a metric of both data quality and data size. In HDX-MS data, a higher average peptide coverage redundancy means more overlap of peptide segments, which provides more data for that region and may allow inference of residue level deuteration uptake properties.^64^ Typically, a given HDX-MS data acquisition will seek to minimize resolution and maximize redundancy. The HDX-MS data for the three proteins in the validation set have an average peptide length of 9.3 residues and an average peptide coverage redundancy of 3.8 peptides per residue (Supplemental Table S11). We investigated the effects of these HDX-MS data quality factors on the performance of MInt-HDX by artificially modulating the average peptide length and average redundancy of the HDX-MS data in the validation set and assessing its effect on MInt-HDX’s accuracy.

To probe the effect of peptide resolution, peptides were separated into two groups: peptides with length greater than 9 residues and peptides with length lesser than 10 residues. The selection criteria resulted in two groups with average peptide length of 13.6 and 6.6 residues, respectively (Supplemental Table S11) and with protein sequence coverages of 69.3% and 70.3% and redundancy of 2.9 and 2.3, respectively (Supplemental Table S11). When using the high peptide length group with MInt-HDX, we observed a significant decrease in accuracy with a Top- 1 L-RMSD of 5.8 Å and a Top-3 L-RMSD of 6.4 Å (Table 2, Supplemental Table S10) compared to the control group (entire dataset, Top-1 L-RMSD 2.4 Å and Top-3 L-RMSD 3.0 Å). However, this was also observed for the low peptide length group, which produced L-RMSD values of 5.6 Å and 6.0 Å for the Top-1 and Top-3 poses, respectively (Table 2, Supplemental Table S10). This observation would suggest that the performance of MInt-HDX is not affected by HDX-MS data resolution since both groups perform poorly with L-RMSD above 5 Å, possibly indicating that other factors may have influenced MInt-HDX’s performance. A more detailed look across the individual proteins in the validation set suggested that the decreased performance of MInt-HDX was potentially correlated with HDX-MS data size (number of peptides) rather than resolution.

**Table 2:** The effect of average peptide length and coverage redundancy on MInt-HDX’s performance. Mint-HDX was performed on reduced data set to reflect reduced HDX-MS data quality factors such as peptide length (>9 Residues vs <10 residues) and redundancy (reduce redundancy to ∼1). All docking was performed on three protein systems of the validation set, for 3 replicates each. Shown are the L-RMSD in (Å) of Top-1 pose and Top-3 poses, averaged across all 9 runs with standard deviations.

|  | Average Top-1 |  | Average Top-3 |  |
| --- | --- | --- | --- | --- |
|  | L-RMSD | SD | L-RMSD | SD |
| Resolution |  |  |  |  |
| >9 Residues | 5.8 | 2.5 | 6.4 | 1.9 |
| <10 Residues | 5.6 | 2.2 | 6.0 | 3.8 |
| Redundancy |  |  |  |  |
| No Redundancy | 7.1 | 2.9 | 5.9 | 2.8 |

Indeed, for VDR, the low peptide length group constituted 75% of all peptides and resulted in greater accuracy with Top-1 L-RMSD of 4.3 Å and Top-3 L-RMSD of 6.0 Å while the high peptide length group constituted 25% of all peptides and resulted in a Top-1 L-RMSD of 7.1 Å and Top-3 L-RMSD of 7.4 Å (Supplemental Table S10 and S11). In contrast, for ERK2, the high peptide length group constituted 55% of all peptides and resulted in greater accuracy with Top-1 L-RMSD of 3.1 Å and Top-3 L-RMSD of 5.0 Å while the low peptide length group constituted 45% of all peptides and resulted in a Top-1 L-RMSD of 3.5 Å and Top-3 L-RMSD of 6.3 Å (Supplemental Table S10 and S11).

To probe the effect of HDX-MS data size, we selected a subset of peptides to set the average coverage redundancy across the validation dataset as low as possible, while striving to keep both the average peptide length and the sequence coverage of the proteins the same as that of the control group. The selected peptides accounted for 32% of the total number of peptides in the control group and resulted in an average redundancy of 1.13 (vs. 3.84 for the control group). In contrast, the average peptide length was 8.6 (vs. 9.3 for the control group) and the average sequence coverage was 90% (vs. 94% for the control group) (Supplemental Table S11). The reduced data set obtained by minimizing the redundancy significantly affected the performance of MInt-HDX, leading to a Top-1 L-RMSD of 7.1 Å and a Top-3 L-RMSD of 5.9 Å (Table 2, Supplemental Table S10). The loss in redundancy rendered MInt-HDX inaccurate for modeling protein-small-molecule interactions, confirming the importance of high volume, higher redundancy HDX-MS data. This suggests that the additional information in redundant peptides allows the models to extract more accurate structural information, improves the performance of MInt-HDX, and increases the likelihood of extracting accurate docking poses.

#### 3.2.5 Overcoming Missing HDX-MS Coverage by Modulating Confidence Threshold

Another critical experimental factor that may affect interpretability is the protein sequence coverage. The protein sequence coverage is the percentage of a protein sequence that is covered by at least one peptide segment. Typically, HDX-MS data acquisition aims to maximize protein sequence coverage with coverages of 90% or more often attainable. Lower coverage is often related to either lower quality data or challenging protein systems. Localized missing sequence coverage can potentially affect the proper performance of MInt-HDX. Indeed, it is conceivable that the model predictions may be adversely affected by missing sequence coverage at or near the ligand binding pocket. To assess the consequences of missing localized sequence coverage, we used an HDX-MS dataset which was held out of the training/validation datasets due to missing coverage near the binding pocket. The protein system is 1-deoxy-D-xylulose 5-phosphate synthase (DXPS), an essential thiamin diphosphate (ThDP)-dependent enzyme at a metabolic branchpoint in bacteria, apicomplexan parasites, and plants. DXPS catalyzes the conversion of pyruvate and D-glyceraldehyde 3-phosphate (D-GAP) to DXP in the first committed step of the essential methylerythritol phosphate (MEP) pathway for isoprenoid biosynthesis (Supplemental Figure 5A). DXP is also a precursor in the biosynthesis of vitamins B1 and B6. As it is absent in humans, this branchpoint enzyme is a promising antimicrobial drug target and is currently the subject of research to develop small-molecule inhibitors. We performed HDX-MS on *Deinococcus radiodurans* DXPS (*Dr*DXPS) in the presence or absence of methyl acetylphosphonate (MAP). MAP is a pyruvate analog that inhibits DXPS by competing with pyruvate for active-site binding and forming a covalent phosphonolactylThDP adduct (DXPS-PLThDP, Supplemental Figure 5A) which mimics the C2α-lactylThDP (LThDP) adduct formed in the canonical DXPS reaction (DXPS-LThDP, Supplemental Figure 5). The inspection of HDX-MS data obtained for *Dr*DXPS in the presence or absence of MAP revealed two primary findings concerning its use as input for MInt-HDX. First, MAP resulted in protection from hydrogen-deuterium exchange in multiple areas consistent with ligand binding (Supplemental Figure S5B and S5C). However, the strongest protection was observed for a region ranging from residues 284 to 321 which is characterized by EX1 kinetics (Supplemental Figure S5D). This region has been previously shown to have EX1 kinetics in *E.coli* DXPS^65^ and is near the MAP binding site, as determined by the analysis of the crystal structure of *Dr*DXPS bound to the PLThDP adduct derived from MAP (Supplemental Figure S5B, PDB: 6OUV).^60^ The EX1 kinetics are significantly slowed down in the presence of MAP as previously observed for *E. coli* DXPS, ^65^ however, the XGBoost models within MInt-HDX have not been trained on data containing EX1 kinetics and must be removed from the peptide list input of MInt-HDX. In addition, while the coverage was relatively high at approximately 88%, there were multiple regions of missing coverage surrounding the binding site (Supplemental Figure 6).

Using this data as input, we applied Mint-HDX to model the interaction between *Dr*DXPS and the PLThDP adduct derived from MAP. Due to the lack of a high-resolution structure of unliganded *Dr*DXPS, AlphaFold III^24^ was used to generate an input structure of *Dr*DXPS (Supplemental Figure S7). Using default parameters (confidence threshold = 0.7, β coefficient = 0.6), MInt-HDX returned poor values compared to the conformation of the PLThDP adduct derived from MAP in 6OUV.pdb with an average Top-1 L-RMSD of 19.4 Å and an average Top-3 L-RMSD of 19.6 Å (Table 3, Supplemental Table S12), suggesting that local missing coverage near the ligand binding site can in fact adversely affect its performance. A closer inspection of the results from individual steps of MInt-HDX’s workflow reveals that the missing coverage at the binding site adversely affects the first step of the workflow, the generation of candidate ligand docking sites. Indeed, two candidate docking sites were generated at a confidence threshold of 0.7 (Fig 3A). The alignment of the 6OUV revealed that the crystallographic MAP molecules were not contained within these candidate sites (Fig 3A), therefore biasing subsequent steps of MInt-HDX. To test this hypothesis and address this limitation, we decreased the confidence threshold parameter to increase the number and volume of candidate docking sites. The comparison between the default confidence threshold value of 0.7 and the modified value of 0.5 can be seen in Figure 3. By decreasing the confidence threshold to 0.5, four candidate sites are generated which now encompass the missing coverage areas as well as the crystallographic ligand conformation (Fig 3B).

**Figure 3:**
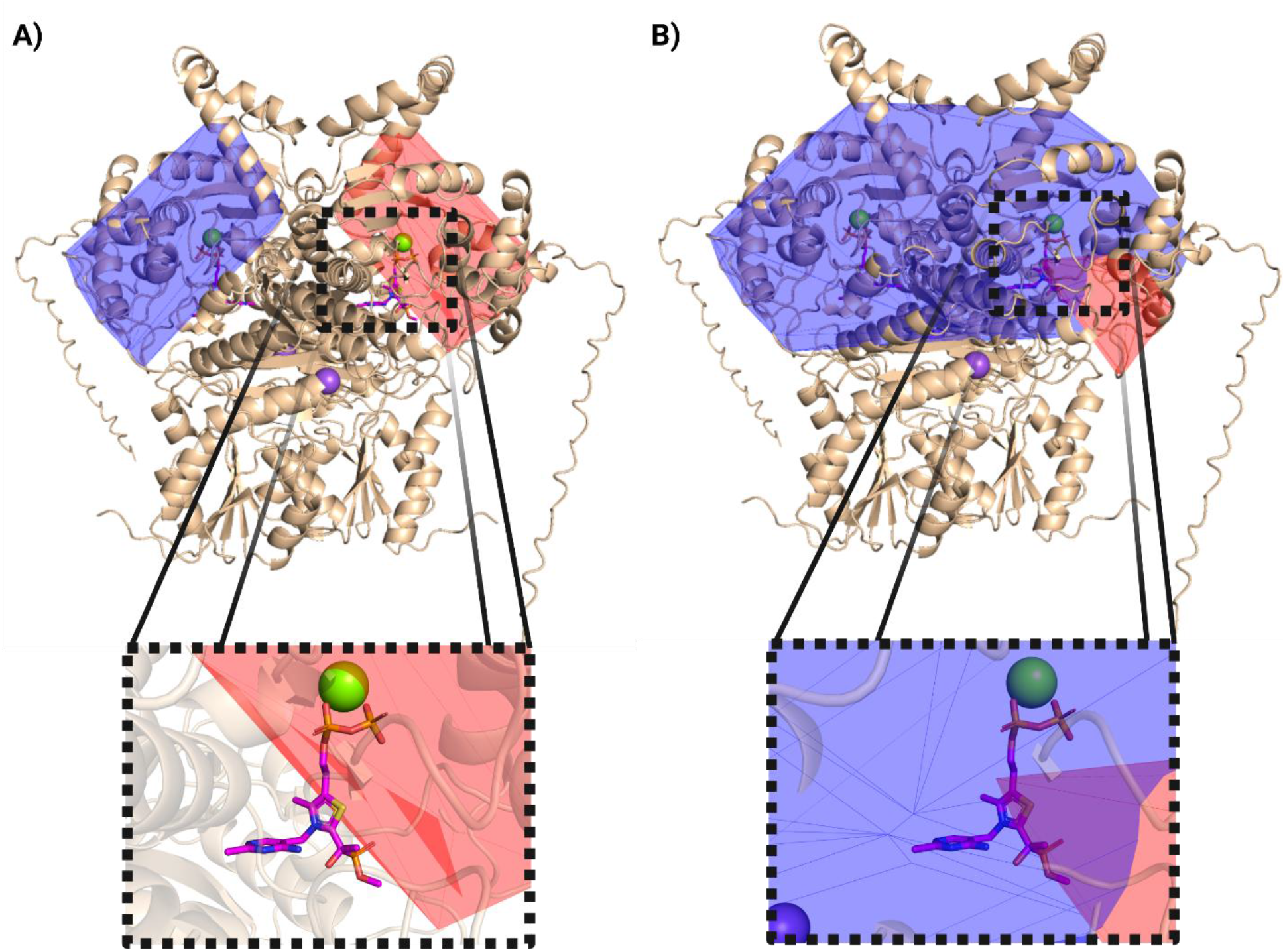
Lowering the confidence threshold to modulate the number and size of candidate docking sites. Candidate docking sites (shaded polygonal 3D shapes) are mapped on the AlphaFold modeled unliganded structure of *Dr*DXPS (cartoon representation) at a confidence threshold parameter of **A)** 0.7, or 70% probability or **B)** 0.5 or 50% probability. The PLThDP adduct derived from MAP was inserted via structural alignment with 6OUV.pdb. Insets show zoomed in view.

**Table 3:** Lowering the confidence threshold improves the performance of MInt-HDX with *Dr*DXPS and overcomes limitations from localized missing sequence coverage. Shown are the average L-RMSD values in Å of the Top-1 and Top-3 poses resulting from MInt-HDX performed on *Dr*DXPS +/- PLThDP adduct derived from MAP at a confidence threshold value of 0.7 and 0.5, respectively. Values were obtained from triplicate runs of MInt-HDX.

|  | Average Top-1 |  | Average Top-3 |  |
| --- | --- | --- | --- | --- |
|  | L-RMSD | SD | L-RMSD | SD |
| Confidence Threshold 0.7 |  |  |  |  |
| MInt-HDX | 19.4 | 0.06 | 19.6 | 0.9 |
| Confidence Threshold 0.5 |  |  |  |  |
| MInt-HDX | 1.7 | 0.01 | 1.8 | 0.2 |

Consequently, when using a confidence threshold of 0.5, MInt-HDX returned vastly improved results. Indeed, as shown in Table 3 and Figure 4, MInt-HDX is able to predict the conformation of the PLThDP adduct derived from MAP in the binding site of *Dr*DXPS at sub 2 Å accuracy with an average Top-1 pose L-RMSD of 1.7 Å and an average Top-3 poses L-RMSD of 1.8 Å compared to the crystallographic conformation (PDC: 6OUV). This demonstrates that modulation of confidence thresholds to expand docking boxes can serve as a strategy to overcome deficiencies in HDX-MS data quality such as localized missing sequence.

**Figure 4:**
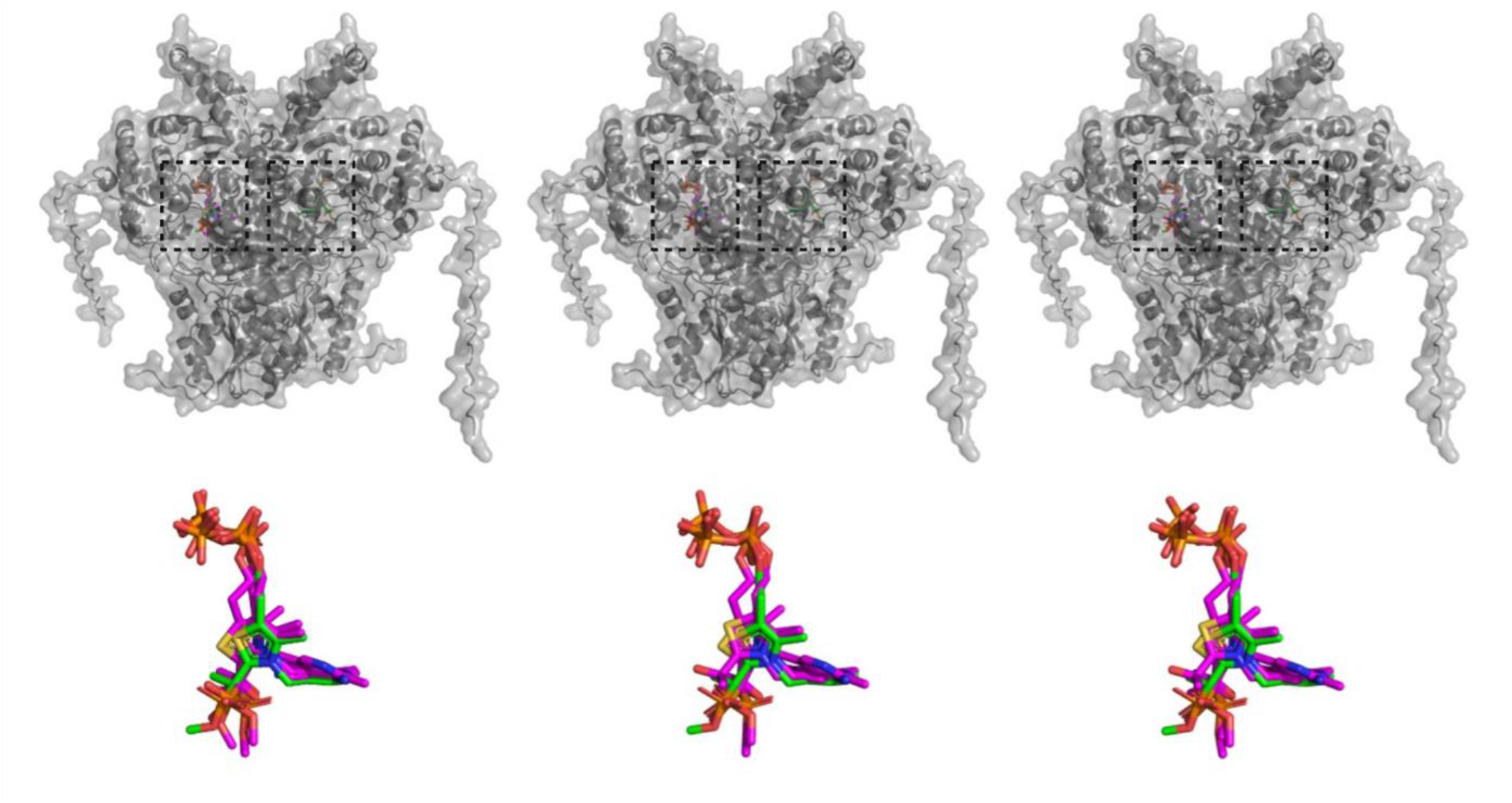
*Dr*DXPS MInt-HDX docking at confidence threshold of 0.5. The docking poses from MInt-HDX (Ligand in Magenta) were aligned with the crystal structure 6OUV.pdb bound to the PLThDP adduct derived from MAP (PLThDP ligand in green). Alignment is performed based on protein backbone atoms. Shown are the Top-3 docking poses from triplicate MInt-HDX runs. The top shows the zoomed out, full protein view with the ligands in sticks and protein in surface representation. The bottom images illustrate the PLThDP ligand conformations zoomed in, without the protein.

#### 3.2.6 MInt-HDX is sensitive enough to identify ligand binding mode in c-src kinase

Finally, we further investigated the sensitivity of MInt-HDX using a special test case, Imatinib binding to c-src kinase. In protein kinases, the conformation of the DFG (Asp-Phe-Gly) motif is closely linked to the activity of the kinase. The DFG-out conformation is considered an inactive state of the kinase, whereas the DFG-in conformation is typically associated with an active state. For ATP competitive inhibitors of protein kinases, the conformation of the DFG motif is also linked to the binding mode of the small-molecule inhibitor which may preferentially target one conformation or the other.^66,67^ Alternatively, the binding of the inhibitor may also be associated with a conformational change of the DFG region.^66,67^

In the case of c-src kinase (the src proto-oncogene, non-receptor tyrosine kinase),^68^ crystallographic evidence (PDB ID 2OIQ) has shown that the conformation of the DFG motif of c- src kinase favors the inactive, DFG-out conformation upon binding to the inhibitor Imatinib (Figure 5A).^69^ However, other crystallographic evidence (PDB ID 1Y57) has shown that MPZ, a des- methyl analog of Imatinib with differing by a single methyl substituent, binds c-src kinase in the active, DFG-in conformation (Figure 5B),^70^ indicating that nearly identical ligands can bind the same target with significantly different conformations (Imatinib vs. MPZ L-RMSD of 12.9 Å ). Thus, we tested whether MInt-HDX is sensitive enough to discern the correct binding mode between c-src kinase and Imatinib. To that end, we applied the MInt-HDX workflow using either the DFG-in or DFG-out conformations as input structures together with the structure of Imatinib and previously acquired HDX-MS data of c-src kinase in the presence and absence of Imatinib.^71^ It is important to note that due to the lack of a ligand-free crystal structure of c-src kinase in a DFG-out conformation, a pseudo-apo DFG-out structure was generated by removing the Imatinib ligand from chain A of 2OIQ. Similarly, a pseudo-apo DFG-in conformation was generated by removing the MPZ ligand from the 1Y57 structure. Finally, chain B of the asymmetric unit of 2OIQ was also used. Electron density for Imatinib is absent in Chain B of 2OIQ and pair-wise all-atom RMSD calculations of the DFG motif suggest that Chain B is closer to a DFG-in conformation (Supplemental Table S13).

**Figure 5:**
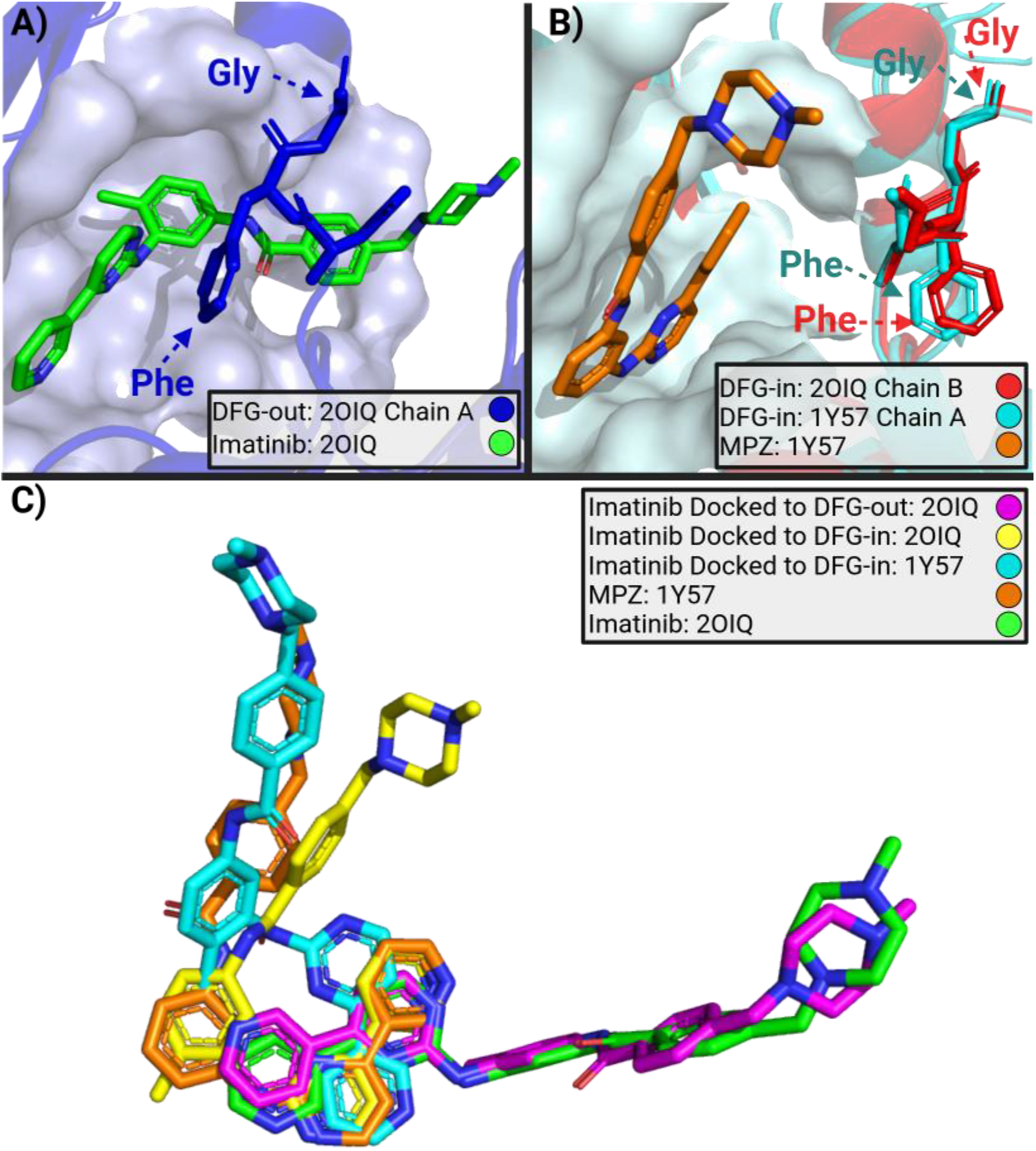
MInt-HDX discerns the DFG conformation of c-src kinase upon Imatinib binding. **A)** C-src kinase in DFG-out conformation (PDB: 2OIQ, Chain A, Blue) upon binding to Imatinib (Green) with the DFG motif shown as sticks. **B)** C-src kinase in DFG-in conformations (PDB: 1Y57, cyan and PDB: 2OIQ, Chain B, red) with MPZ, a des-methyl analog of Imatinib (PDB: 1Y57, orange), and the DFG motif shown as sticks. **C)** Alignment of the top docking pose of Imatinib resulting from MInt-HDX, when docked to different conformations of c-src kinase as follows: docked to the DFG-out 2OIQ chain A (magenta), docked to the DFG-in 2OIQ Chain B (yellow) and docked to the DFG-in 1Y57 (cyan). Imatinib from 2OIQ chain A is shown in green and MPZ from 1Y57 is shown in orange. Structural alignment was performed based on protein backbone atoms.

Looking at the resulting Top-1 pose, MInt-HDX results were heavily dependent on the DFG conformation. Indeed, as shown visually in Figure 5 Panel C and quantitatively from the L-RMSD values in Supplemental Table S13, when performing MInt-HDX with an input structure in a DFG-in conformation (1Y57 or Chain B of 2OIQ), the resulting top poses were more consistent with the MPZ conformation in 1Y57. On the other hand, when using the DFG-out conformation as an input structure (Chain A of 1OIQ), the top pose was more consistent with Imatinib conformation in 2OIQ chain A. While it is not surprising that the docking results depend on the conformation of the binding pocket in the input structure, we can further assess which of these top poses is most consistent with the HDX-MS data by examining their Score_Avg_, the scoring function outputs of MInt-HDX. The highest-scored overall pose across the three input conformations was determined to be that resulting from the DFG-out input structure 2OIQ-ChainA, with the top pose having an average score of 5.6 across the three replicates(Supplemental Table S13). On the other hand, the top poses of the DFG-in inputs using 2OIQ-chain B or 1Y57 had a lower average score of 4.0 (Supplemental Table S13). These results are in agreement with the consensus that Imatinib favors the DFG-out conformation,^66^ however, it does not rule out the existence of a minor population of Imatinib bound to the protein in a DFG-in conformation as part of a complex binding landscape.

These results demonstrate that the MInt-HDX workflow is sensitive enough to discern and provide insights into binding modes within a binding pocket if provided with an adequate range of input structures that adequately represents the conformational landscape of the binding pocket. Currently, MInt-HDX takes advantage of a rigid docking method within its workflow. Future work will involve incorporating flexible docking approaches to model ligand binding that will account for conformational heterogeneity of the target protein to address induced fit, conformational selection or co-folding processes.

## 4 Conclusion

In this work, we developed MInt-HDX, a hybrid physics-based/ML-based framework that leverages HDX-MS data to accurately model protein-small-molecule interactions, demonstrating the value of integrating lower resolution experimental data with modern computational approaches. By combining the mechanistic and biophysical foundation of physics-based molecular docking approaches and the pattern recognition capabilities of machine learning with peptide-level resolution HDX-MS, MInt-HDX achieves sub-3 Å ligand pose prediction accuracy, a level suitable for structure-based drug discovery efforts and/or medicinal chemistry applications. These results establish MInt-HDX as a reliable tool for modeling biologically and pharmacologically relevant binding modes. Notably, MInt-HDX’s hybrid workflow consistently outperforms conventional molecular docking approaches lacking *a priori* binding information by instead integrating HDX-MS data interpreted by HDX-MS-trained XGBoost models. Furthermore, its performance and accuracy is comparable to that of ML docking methods, despite being trained on vastly smaller training datasets. This demonstrates that incorporating experimentally derived HDX-MS constraints and physics-based approaches together with ML-based and artificial intelligence frameworks can compensate for the need for massive training datasets and provide more data-efficient avenues toward biologically and pharmacologically accurate structural prediction. MInt-HDX also displays comparable accuracy to our previously reported^35^ HDX-MS-guided physics-based modeling approach which employs maximum entropy-based ensemble reweighting. However, it vastly improves efficiency with reduced computational costs and turnaround time. Rather than requiring extensive molecular dynamics simulations, *post hoc* ensemble reweighting procedures and dimensional reduction techniques to extract docking poses that may require days to weeks of computation, MInt-HDX produces predictions within minutes to hours with comparable structural accuracy. This improved efficiency significantly enhances the feasibility of incorporating HDX-MS-guided structural modeling into iterative drug discovery workflows, where rapid hypothesis generation and evaluation are essential for lead optimization and compound prioritization.

Beyond the application to protein-ligand docking and drug discovery, MInt-HDX’s performance also highlights the value of HDX-MS as an information-rich structural biology technique ideal for integrating with computational approaches. Although HDX-MS does not directly provide atomic resolution information, it reports on detailed information regarding localized biophysical properties such as local protein dynamics and conformational equilibria, solvent accessibility, hydrogen bonding, and allosteric propagation.^29,30,64^ Expansion of hybrid integrative strategies such as MInt-HDX would enable the extraction of increasingly rich structural, dynamic and mechanistic information from HDX-MS data while preserving physical interpretability and experimental grounding. As these computational paradigms continue to evolve, HDX-MS can be an important source of experimental supervision for modeling protein systems whose intrinsic conformational heterogeneity remains challenging to characterize by traditional structural biology methods. Future developments will focus on extending this framework to encompass flexible ligand docking, and then beyond protein-small-molecule interactions to protein–protein and protein–nucleic acid interactions. In parallel, we envision expanding the methodology to modeling protein conformational ensembles by integrating HDX-MS with enhanced sampling techniques and next-generation machine learning models to characterize the distribution of biologically relevant conformations rather than a single static structure. Such developments have the potential to establish HDX-MS-guided hybrid computational modeling as a valuable platform for studying biomolecular recognition, allostery, and conformational landscapes.

## Supporting Information

Additional parameters, data and results referenced in the main text (PDF)

## Acknowledgments

Abstract Figure *Created in BioRender. Lowe, V. (2026)* https://BioRender.com/fmmq1gg

## Data Sharing Statement

### Data Availability

Data available upon contacting corresponding author at

## Code Availability

Contact corresponding author at. https://github.com/Deredge-Lab-UMB/HDXpert-Dock

## Transparency Declaration

A patent is currently pending for the work in the manuscript.

## Terminology and Abbreviation

eXtreme Gradient Boosting: XGBoost
XGBoost Models: Inter-XGB, Dist-XGB, SASA-XGB
Density Based Spatial Cluster of Applications with Noise: DBSCAN
Ligand RMSD: L-RMSD
Deuteration Difference: ΔD
Docking Predictions of Target Values: Inter-Dock, Dist-Dock, SASA-Dock
Hydrogen-Deuterium Exchange Mass Spectrometry: HDX-MS
Modeling by Integrative HDX-MS: Mint-HDX
Hydrogen-Deuterium Exchange Ensemble Reweighting: HDXer
*Deinococcus radiodurans* 1-deoxy-D-xylulose 5-phosphate synthase: *Dr*DXPS
Methyl acetylphosphonate: MAP

