## Supplemental Information for "MInt-HDX: Leveraging Hydrogen-Deuterium Exchange Mass Spectrometry and Machine-Learning to Improve Protein-Ligand Docking"

**Table of Contents:**


| | |
|---|---|
| Supplemental Table S1: Training set proteins | Page 3 |
| Supplemental Table S2: Validation set proteins | Page 3 |
| Supplemental Table S3: RMSD of ligand-bound vs unbound crystal structures | Page 4 |
| Supplemental Table S4: Inter-XGB model hyperparameters and performance metrics | Page 4 |
| Supplemental Table S5: Dist-XGB model hyperparameters and performance metrics | Page 5 |
| Supplemental Table S6: SASA-XGB model hyperparameters and performance metrics | Page 5 |
| Supplemental Table S7: The effect of confidence threshold on candidate docking sites generation | Page 6 |
| Supplemental Table S8: Filtering function and optimization of β coefficient | Page 7 |
| Supplemental Table S9: Comparison of scoring functions across all replicas | Page 8 |
| Supplemental Table S10: Validation set L-RMSD across replicas and methods | Page 9 |
| Supplemental Table S11: HDX-MS peptide segment length, redundancy, and coverage | Page 10 |
| Supplemental Table S12: L-RMSD of replicas of MAP docking to DrDXPS at confidence threshold 0.5 and 0.7 | Page 11 |
| Supplemental Table S13: C-src kinase replica MInt-HDX docking, L-RMSD and scores, and crystal structure RMSD's of the DFG region and ligand | Page 12 |
| Supplemental Figure 1: XGBoost models performance on the validation set data | Page 12 |
| Supplemental Figure 2: Three docking replicates of each docking tool to VDR | Page 13 |
| Supplemental Figure 3: Three docking replicates of each tool to ERK2 | Page 14 |
| Supplemental Figure 4: Three docking replicates of each tool to FOSA | Page 15 |
| Supplemental Figure 5: HDX-MS data for DrDXPS | Page 16 |
| Supplemental Figure 6: Areas of missing HDX-MS coverage mapped to the crystal structure of DrDXPS | Page 17 |
| Supplemental Figure 7: AlphaFold III model confidence map for the apo form of DrDXPS | Page 17 |

| PRIDE ID / In House | System | Ligand | Species | Classification | Size (Residues) | PDB | Time Points (Min) | Instrument | pH | Conditions |
|---|---|---|---|---|---|---|---|---|---|---|
| PXD037721 | PI3Kα | NVP-BYL719 | *Homo sapiens* | Kinase | 1355 | 7PG6 | 0.05, 0.5, 5, 50 | Synapt G2-Si | 7.5 | 150 mM NaCl |
| PXD037721 | PI3Kα | 1938 | *Homo sapiens* | Kinase | 946 | 8OW2 | 0.05, 0.5, 5, 50 | Synapt G2-Si | 7.5 | 150 mM NaCl |
| PXD037448<br>PXD037374<br>PXD037355 | eGFR Exon | Erlotinib | *Homo sapiens* | Kinase | 333 | 1M17 | 0.167, 1, 3, 10 | Synapt G2-Si | 7 | 100 mM NaCl |
| PXD037151 | NicA2 | N-methylmyosmine | *Pseudomonas putida* | Oxidoreductase | 864 | 7C4A | 0.5, 2, 5 | Synapt G2-Si | 7.6 | 10 mM potassium phosphate |
| In House | HasA | Heme | *Pseudomonas aeruginosa* | Hemophore | 205 | 3ELL | 0.17, 1, 10, 60, 120, 500 | Synapt G2-Si | 7.4 | 20 mM sodium phosphate buffer |
| In House | HasA | Salophen | *Pseudomonas aeruginosa* | Hemophore | 205 | 3W8M | 0.17, 1, 10, 60, 120, 500 | Synapt G2-Si | 7.4 | 20 mM sodium phosphate buffer |
| In House | HCV | Filibuvir | Hepatitis C Virus | Genome polyprotein / Transferase | 562 | 3FRZ | 0, 0.17, 10, 60, 120, 500 | Synapt G2-Si | 7.4 | 150 mm NaCl, 5 mm DTT, 2 mm $MgCl_2$ |
| In House | ERK2_UP | Tetrahydropyrrolo-diazepenone (BVD) | *Homo sapiens* | Kinase | 330 | 5BVD | 0, 0.17, 1, 10, 60, 120, 500 | Synapt G2-Si | 7.4 | 300 mM NaCl |
| In House | ERK2_UP | GDC-0994 | *Homo sapiens* | Kinase | 348 | 6OPH | 0, 0.17, 1, 10, 60, 120, 500 | Synapt G2-Si | 7.4 | 300 mM NaCl |
| In House | ERK2_UP | SCH772984 | *Homo sapiens* | Kinase | 335 | 4QTA | 0, 0.17, 1, 10, 60, 120, 500 | Synapt G2-Si | 7.4 | 300 mM NaCl |
| In House | ERK2_UP | VTX-11e | *Homo sapiens* | Kinase | 348 | 4QTE | 0, 0.17, 1, 10, 60, 120, 500 | Synapt G2-Si | 7.4 | 300 mM NaCl |

**Supplemental Table S1: Training set proteins.** All proteins included in the training set of MInt-HDX. For each protein, the PRIDE ID is listed if applicable, along with information on the system, ligand, species, and other structural factors.

| PRIDE ID / In House | System | Ligand | Species | Classification | Size (Residues) | PDB | Time Points (Min) | Instrument | pH | Conditions |
|---|---|---|---|---|---|---|---|---|---|---|
| PXD019810 | VDR | Vitamin D (VDX) | *Homo sapiens* | Vitamin D Receptor | 250 | 1DB1 | 0.5, 1, 3, 10, 30 | Synapt G2-Si | 8 | 200 mM NaCl |
| PXD048311 | ERK2_UP | Vertex-11e | *Rattus norvegicus* | Kinase | 346 | 6OPK | 0.5, 1, 3, 8, 16, 30, 60, 90, 180 | Synapt G2 HDMS Q-TOF | 7.2 | 100 mM NaCl, 5 mM DTT, 10 mM $MgCl_2$ |
| In House | FosA | ANY1 | *Escherichia coli* | Metalloenzymes/Transferase | 288 | 5WEW | 0, 0.17, 1, 10, 60, 120, 500 | Synapt G2-Si | 7.8 | 150 mM Kcl, 50 µM MnCl2 |
| In House | DXPS | MAP | *Deinococcus radiodurans* | Synthase | 1171 | 6OUV | 0, 0.17, 0.33, 1, 5, 10, 60, 120 | Synapt G2-Si | 8 | 50 mM NaCl, 1 mM MgCl2 |

**Supplemental Table S2: Validation set proteins.** All proteins included in the validation set of MInt-HDX. For each protein, the PRIDE ID is listed if applicable, along with information on the system, ligand, species, and other structural factors.

| System | Ligand | RMSD |
|---|---|---|
| PI3Kα | NVP-BYL719 | 0.7 |
| PI3Kα | 1938 | 2.1 |
| eGFR Exon | Erlotinib | 1.1 |
| NicA2 | N-methylmyosmine | 26.6 |
| HasA | Heme | 5.6 |
| HasA | Salophen | 5.5 |
| HCV | Filibuvir | 1 |
| ERK2_UP | Tetrahydropyrrolo-diazepenone (BVD) | 1.9 |
| ERK2_UP | GDC-0994 | 1.3 |
| ERK2_UP | SCH772984 | 1.9 |
| ERK2_UP | VTX-11e | 1.5 |
| VDR | Vitamin D (VDX) | 0.5 |
| ERK2_UP | Vertex-11e | 1.2 |
| FosA | ANY1 | 1.1 |

**Supplemental Table S3: RMSD of ligand-bound vs unbound crystal structures.** The RMSD difference between the ligand-bound and unbound crystal structures for each protein in the training and validation set.

| Parameter | Search Space |
|---|---|
| Lambda | Float Value from 0 to 5, **0** |
| Gamma | Float Value from 0 to 5, **3.36** |
| Max Depth | Integer Value from 3 to 12, **5** |
| Learning Rate | **0.001** |
| Min Child Depth | Integer Value from 1 to 250, **3** |
| Column Sampling by Node | Float Value from 0.1 to 1, **0.79** |
| Subsample | Float Value from 0.1 to 1, **0.84** |
| Number of Boost Rounds | **1429** |
| Early Stopping Rounds | **100** |
| Tree Method | Approximate, Hist, **Exact** |
| Precision [Class 0, Class 1, Weighted Average] | [0.88, 0.58, 0.78] |
| Recall [Class 0, Class 1, Weighted Average] | [0.72, 0.80, 0.74] |
| F1 Score [Class 0, Class 1, Weighted Average] | [0.79, 0.67, 0.75] |

**Supplemental Table S4: Inter-XGB model hyperparameters and performance metrics.** The hyperparameters for the Inter-XGB model that were searched through, with the final parameter used in the model in bold. Additionally, the performance metrics of precision, recall, and F1 Score for the model by class and aggregated by the weighted average are shown to indicate model performance

| Parameter | Search Space |
| --- | --- |
| Lambda | Float Value from 0 to 5, **2.33** |
| Gamma | Float Value from 0 to 5, **0.86** |
| Max Depth | Integer Value from 3 to 12, **8** |
| Learning Rate | **0.001** |
| Min Child Depth | Integer Value from 1 to 250, **8** |
| Column Sampling by Node | Float Value from 0.1 to 1, **0.73** |
| Subsample | Float Value from 0.1 to 1, **0.85** |
| Number of Boost Rounds | **925** |
| Early Stopping Rounds | **100** |
| Tree Method | Approximate, Hist, **Exact** |
| Precision [Class 0, Class 1, Weighted Average] | [0.71, 0.66, 0.69] |
| Recall [Class 0, Class 1, Weighted Average] | [0.72, 0.65, 0.69] |
| F1 Score [Class 0, Class 1, Weighted Average] | [0.71, 0.66, 0.69] |

**Supplemental Table S5: Dist-XGB model hyperparameters and performance metrics.** The hyperparameters for the Dist-XGB model that were searched through, with the final parameter used in the model in bold. Additionally, the performance metrics of precision, recall, and F1 Score for the model by class and aggregated by the weighted average are shown to indicate model performance

| Parameter | Search Space |
| --- | --- |
| Lambda | Float Value from 0 to 5, **0.09** |
| Gamma | Float Value from 0 to 5, **3.32** |
| Max Depth | Integer Value from 3 to 12, **3** |
| Learning Rate | **0.001** |
| Min Child Depth | Integer Value from 1 to 250, **1** |
| Column Sampling by Node | Float Value from 0.1 to 1, **0.54** |
| Subsample | Float Value from 0.1 to 1, **0.74** |
| Number of Boost Rounds | **1623** |
| Early Stopping Rounds | **100** |
| Tree Method | Approximate, Hist, **Exact** |
| Precision [Class 0, Class 1, Weighted Average] | [0.92, 0.57, 0.82] |
| Recall [Class 0, Class 1, Weighted Average] | [0.72, 0.86, 0.76] |
| F1 Score [Class 0, Class 1, Weighted Average] | [0.81, 0.69, 0.77] |

**Supplemental Table S6: SASA-XGB model hyperparameters and performance metrics.** The hyperparameters for the SASA-XGB model that were searched through, with the final parameter used in the model in bold. Additionally, the performance metrics of precision, recall, and F1 Score for the model by class and aggregated by the weighted average are shown to indicate model performance.

| System | Confidence Threshold | Number of Candidate Docking Sites | Ligand Within Docking Box | Candidate Docking Site 1 Volume (Å³) | Candidate Docking Site 2 Volume (Å³) | Candidate Docking Site 3 Volume (Å³) |
|---|---|---|---|---|---|---|
| ERK2 | 0.5 | 1 | Y | 25639 | 0 | 0 |
| ERK2 | 0.6 | 1 | Y | 20764 | 0 | 0 |
| ERK2 | 0.7 | 2 | Y | 2177 | 1827 | 0 |
| ERK2 | 0.8 | 0 | N | 0 | 0 | 0 |
| VDR | 0.5 | 1 | Y | 20770 | 0 | 0 |
| VDR | 0.6 | 1 | Y | 16240 | 0 | 0 |
| VDR | 0.7 | 1 | Y | 13559 | 0 | 0 |
| VDR | 0.8 | 0 | N | 0 | 0 | 0 |
| FosA | 0.5 | 1 | Y | 20458 | 0 | 0 |
| FosA | 0.6 | 3 | Y | 3896 | 1242 | 1307 |
| FosA | 0.7 | 2 | Y | 1242 | 1307 | 0 |
| FosA | 0.8 | 0 | N | 0 | 0 | 0 |

**Supplemental Table S7: The effect of confidence threshold on candidate docking sites generation.** For each protein in the validation set, the confidence threshold value was scanned from 0.5 to 0.8, and the number of candidate docking sites, the presence of the ligand within the docking box, and the candidate docking site volume (in Å$^3$) are shown in the table.

| Protein | Replicate | β | > 10 Å, Kept (FP) | < 10 Å, Removed (FN) | < 10 Å, Kept (TP) | > 10 Å, Removed (TN) |
|---|---|---|---|---|---|---|
| ERK2 | Rep 1 | 0.1 | 23.0 | 0.0 | 17.0 | 0.0 |
| | | 0.2 | 23.0 | 0.0 | 17.0 | 0.0 |
| | | 0.3 | 23.0 | 0.0 | 17.0 | 0.0 |
| | | 0.4 | 3.0 | 0.0 | 17.0 | 20.0 |
| | | 0.5 | 3.0 | 0.0 | 17.0 | 20.0 |
| | | 0.6 | 3.0 | 0.0 | 17.0 | 20.0 |
| | | 0.7 | 0.0 | 20.0 | 0.0 | 20.0 |
| | | 0.8 | 0.0 | 20.0 | 0.0 | 20.0 |
| | | 0.9 | 0.0 | 20.0 | 0.0 | 20.0 |
| | | 1.0 | 0.0 | 20.0 | 0.0 | 20.0 |
| | Rep 2 | 0.1 | 23.0 | 0.0 | 17.0 | 0.0 |
| | | 0.2 | 23.0 | 0.0 | 17.0 | 0.0 |
| | | 0.3 | 23.0 | 0.0 | 17.0 | 0.0 |
| | | 0.4 | 3.0 | 0.0 | 17.0 | 20.0 |
| | | 0.5 | 3.0 | 0.0 | 17.0 | 20.0 |
| | | 0.6 | 3.0 | 0.0 | 17.0 | 20.0 |
| | | 0.7 | 0.0 | 20.0 | 0.0 | 20.0 |
| | | 0.8 | 0.0 | 20.0 | 0.0 | 20.0 |
| | | 0.9 | 0.0 | 20.0 | 0.0 | 20.0 |
| | | 1.0 | 0.0 | 20.0 | 0.0 | 20.0 |
| | Rep 3 | 0.1 | 24.0 | 0.0 | 16.0 | 0.0 |
| | | 0.2 | 24.0 | 0.0 | 16.0 | 0.0 |
| | | 0.3 | 24.0 | 0.0 | 16.0 | 0.0 |
| | | 0.4 | 4.0 | 0.0 | 16.0 | 20.0 |
| | | 0.5 | 4.0 | 0.0 | 16.0 | 20.0 |
| | | 0.6 | 4.0 | 0.0 | 16.0 | 20.0 |
| | | 0.7 | 0.0 | 20.0 | 0.0 | 20.0 |
| | | 0.8 | 0.0 | 20.0 | 0.0 | 20.0 |
| | | 0.9 | 0.0 | 20.0 | 0.0 | 20.0 |
| | | 1.0 | 0.0 | 20.0 | 0.0 | 20.0 |
| VDR | Rep 1 | 0.1 | 0.0 | 0.0 | 18.0 | 0.0 |
| | | 0.2 | 0.0 | 0.0 | 18.0 | 0.0 |
| | | 0.3 | 0.0 | 0.0 | 18.0 | 0.0 |
| | | 0.4 | 0.0 | 0.0 | 18.0 | 0.0 |
| | | 0.5 | 0.0 | 0.0 | 18.0 | 0.0 |
| | | 0.6 | 0.0 | 0.0 | 18.0 | 0.0 |
| | | 0.7 | 0.0 | 6.0 | 12.0 | 0.0 |
| | | 0.8 | 0.0 | 18.0 | 0.0 | 0.0 |
| | | 0.9 | 0.0 | 18.0 | 0.0 | 0.0 |
| | | 1.0 | 0.0 | 18.0 | 0.0 | 0.0 |
| | Rep 2 | 0.1 | 0.0 | 0.0 | 20.0 | 0.0 |
| | | 0.2 | 0.0 | 0.0 | 20.0 | 0.0 |
| | | 0.3 | 0.0 | 0.0 | 20.0 | 0.0 |
| | | 0.4 | 0.0 | 0.0 | 20.0 | 0.0 |
| | | 0.5 | 0.0 | 0.0 | 20.0 | 0.0 |
| | | 0.6 | 0.0 | 0.0 | 20.0 | 0.0 |
| | | 0.7 | 0.0 | 5.0 | 15.0 | 0.0 |
| | | 0.8 | 0.0 | 18.0 | 2.0 | 0.0 |
| | | 0.9 | 0.0 | 20.0 | 0.0 | 0.0 |
| | | 1.0 | 0.0 | 20.0 | 0.0 | 0.0 |
| | Rep 3 | 0.1 | 0.0 | 0.0 | 19.0 | 0.0 |
| | | 0.2 | 0.0 | 0.0 | 19.0 | 0.0 |
| | | 0.3 | 0.0 | 0.0 | 19.0 | 0.0 |
| | | 0.4 | 0.0 | 0.0 | 19.0 | 0.0 |
| | | 0.5 | 0.0 | 0.0 | 19.0 | 0.0 |
| | | 0.6 | 0.0 | 0.0 | 19.0 | 0.0 |
| | | 0.7 | 0.0 | 6.0 | 13.0 | 0.0 |
| | | 0.8 | 0.0 | 16.0 | 3.0 | 0.0 |
| | | 0.9 | 0.0 | 19.0 | 0.0 | 0.0 |
| | | 1.0 | 0.0 | 19.0 | 0.0 | 0.0 |
| FOSA | Rep 1 | 0.1 | 8.0 | 0.0 | 32.0 | 0.0 |
| | | 0.2 | 8.0 | 0.0 | 32.0 | 0.0 |
| | | 0.3 | 8.0 | 0.0 | 32.0 | 0.0 |
| | | 0.4 | 8.0 | 0.0 | 32.0 | 0.0 |
| | | 0.5 | 8.0 | 0.0 | 32.0 | 0.0 |
| | | 0.6 | 5.0 | 1.0 | 31.0 | 3.0 |
| | | 0.7 | 4.0 | 1.0 | 31.0 | 4.0 |
| | | 0.8 | 3.0 | 16.0 | 16.0 | 5.0 |
| | | 0.9 | 0.0 | 24.0 | 8.0 | 8.0 |
| | | 1.0 | 0.0 | 32.0 | 0.0 | 8.0 |
| | Rep 2 | 0.1 | 8.0 | 0.0 | 32.0 | 0.0 |
| | | 0.2 | 8.0 | 0.0 | 32.0 | 0.0 |
| | | 0.3 | 8.0 | 0.0 | 32.0 | 0.0 |
| | | 0.4 | 8.0 | 0.0 | 32.0 | 0.0 |
| | | 0.5 | 8.0 | 0.0 | 32.0 | 0.0 |
| | | 0.6 | 4.0 | 0.0 | 32.0 | 4.0 |
| | | 0.7 | 4.0 | 1.0 | 31.0 | 4.0 |
| | | 0.8 | 3.0 | 17.0 | 15.0 | 5.0 |
| | | 0.9 | 0.0 | 25.0 | 7.0 | 8.0 |
| | | 1.0 | 0.0 | 32.0 | 0.0 | 8.0 |
| | Rep 3 | 0.1 | 7.0 | 0.0 | 32.0 | 0.0 |
| | | 0.2 | 7.0 | 0.0 | 32.0 | 0.0 |
| | | 0.3 | 7.0 | 0.0 | 32.0 | 0.0 |
| | | 0.4 | 7.0 | 0.0 | 32.0 | 0.0 |
| | | 0.5 | 7.0 | 0.0 | 32.0 | 0.0 |
| | | 0.6 | 3.0 | 0.0 | 32.0 | 4.0 |
| | | 0.7 | 3.0 | 1.0 | 31.0 | 4.0 |
| | | 0.8 | 2.0 | 15.0 | 18.0 | 5.0 |
| | | 0.9 | 0.0 | 26.0 | 7.0 | 7.0 |
| | | 1.0 | 0.0 | 32.0 | 0.0 | 7.0 |

| β | > 10 Å, Kept (FP) | < 10 Å, Removed (FN) | < 10 Å, Kept (TP) | > 10 Å, Removed (TN) | Percent < 10 Å, Kept | Percent > 10 Å, Removed |
|---|---|---|---|---|---|---|
| 0.1 | 93 | 0 | 203 | 0 | 100.00% | 0.00% |
| 0.2 | 93 | 0 | 203 | 0 | 100.00% | 0.00% |
| 0.3 | 93 | 0 | 203 | 0 | 100.00% | 0.00% |
| 0.4 | 33 | 0 | 203 | 60 | 100.00% | 64.52% |
| 0.5 | 33 | 0 | 203 | 60 | 100.00% | 64.52% |
| 0.6 | 22 | 1 | 202 | 71 | 99.51% | 76.34% |
| 0.7 | 11 | 80 | 133 | 72 | 62.44% | 86.75% |
| 0.8 | 8 | 160 | 54 | 75 | 25.23% | 90.36% |
| 0.9 | 0 | 192 | 22 | 83 | 10.28% | 100.00% |
| 1 | 0 | 213 | 0 | 83 | 0.00% | 100.00% |

**Supplemental Table S8: Filtering function and optimization of β coefficient.** β values ranging from 0.1 to 1 were used to filter docking poses for each validation set protein and replicate and analyzed against the filtering criteria of above or below 10 Å. Bottom table shows the average for each β value performance taken across all systems and replicates.

| System | Replicate | | MInt-HDX (Filtered Poses) | AutoDock Vina (Filtered Poses) |
|---|---|---|---|---|
| VDR | Rep 1 | Pose 1 | 2.6 | 0.8 |
| | | Pose 2 | 2.6 | 1.3 |
| | | Pose 3 | 1.3 | 9.2 |
| | | Average | 2.2 | 3.8 |
| | | Diff MInt-HDX | N/A | -1.6 |
| | Rep 2 | Pose 1 | 1.7 | 0.9 |
| | | Pose 2 | 1.9 | 1.3 |
| | | Pose 3 | 0.9 | 1.7 |
| | | Average | 1.5 | 1.3 |
| | | Diff MInt-HDX | N/A | 0.2 |
| | Rep 3 | Pose 1 | 1.7 | 0.9 |
| | | Pose 2 | 3.1 | 1.4 |
| | | Pose 3 | 0.9 | 1.7 |
| | | Average | 1.9 | 1.3 |
| | | Diff MInt-HDX | N/A | 0.6 |
| | **Average Top Pose** | | 2.0 | 0.9 |
| | **Standard Deviation** | | **0.8** | **2.7** |
| | **Standard Deviation (Top-1)** | | **0.5** | **0.0** |
| | **Average Top 3 Poses** | | **1.8** | **2.1** |
| | **Average** | **Difference to MInt-HDX Top 3** | | **-0.3** |
| | | **Difference to MInt-HDX Top 1** | | **1.1** |
| ERK2 | Rep 1 | Pose 1 | 3.1 | 3.1 |
| | | Pose 2 | 4.4 | 6.0 |
| | | Pose 3 | 4.1 | 8.2 |
| | | Average | 3.9 | 5.8 |
| | | Diff MInt-HDX | N/A | -1.9 |
| | Rep 2 | Pose 1 | 3.1 | 3.1 |
| | | Pose 2 | 4.3 | 6.0 |
| | | Pose 3 | 4.7 | 8.2 |
| | | Average | 4.0 | 5.8 |
| | | Diff MInt-HDX | N/A | -1.7 |
| | Rep 3 | Pose 1 | 3.1 | 3.1 |
| | | Pose 2 | 3.3 | 8.2 |
| | | Pose 3 | 4.8 | 6.0 |
| | | Average | 3.7 | 5.8 |
| | | Diff MInt-HDX | N/A | -1.9 |
| | **Average Top Pose** | | **3.1** | **3.1** |
| | **Standard Deviation** | | **0.7** | **2.2** |
| | **Standard Deviation (Top-1)** | | **0.0** | **0.0** |
| | **Average Top 3 Poses** | | **3.9** | **5.8** |
| | **Average** | **Difference to MInt-HDX Top 3** | | **-1.9** |
| | | **Difference to MInt-HDX Top 1** | | **0.0** |
| FosA | Rep 1 | Pose 1 | 2.1 | 2.1 |
| | | Pose 2 | 3.1 | 11.4 |
| | | Pose 3 | 4.0 | 3.4 |
| | | Average | 3.1 | 5.7 |
| | | Diff MInt-HDX | N/A | -2.6 |
| | Rep 2 | Pose 1 | 2.1 | 2.1 |
| | | Pose 2 | 3.8 | 11.4 |
| | | Pose 3 | 3.9 | 3.4 |
| | | Average | 3.3 | 5.6 |
| | | Diff MInt-HDX | N/A | -2.4 |
| | Rep 3 | Pose 1 | 2.1 | 11.4 |
| | | Pose 2 | 3.6 | 3.4 |
| | | Pose 3 | 4.1 | 11.3 |
| | | Average | 3.3 | 8.7 |
| | | Diff MInt-HDX | N/A | -5.6 |
| | **Average Top Pose** | | **2.1** | **5.2** |
| | **Standard Deviation** | | **0.9** | **4.5** |
| | **Standard Deviation (Top-1)** | | **0.0** | **5.4** |
| | **Average Top 3 Poses** | | **3.2** | **6.7** |
| | **Average** | **Difference to MInt-HDX Top 3** | | **-3.5** |
| | | **Difference to MInt-HDX Top 1** | | **-3.1** |
| | **Overall Average** | | **3.0** | **4.9** |
| | **Standard Deviation** | | **0.8** | **3.1** |
| | **Top 3 Pose Diff Avg** | | **N/A** | **-1.9** |
| | **Top Pose Overall Avg** | | **2.4** | **3.1** |
| | **Standard Deviation (Top-1)** | | **0.6** | **2.5** |
| | **Top Pose Diff Avg** | | **N/A** | **-0.7** |

**Supplemental Table S9: Comparison of scoring functions across all replicas.** The L-RMSD of Top-1 and Top-3 docking poses for each validation set protein and replicate were calculated for the MInt-HDX's scoring function and AutoDock Vina's scoring function on the filtered poses.

| System | Replicate | | MInt-HDX | Vina | Swissdock | DiffDock | Peptides (>9 Residues) | Peptides (<10 Residues) | Redundancy |
|---|---|---|---|---|---|---|---|---|---|
| VDR | Rep 1 | Pose 1 | 2.6 | 9.9 | 2.1 | 0.6 | 10.0 | 9.6 | 9.2 |
| | | Pose 2 | 2.6 | 2.7 | 0.9 | 0.6 | 10.0 | 9.7 | 1.3 |
| | | Pose 3 | 1.3 | 2.9 | 9.2 | 0.5 | 2.7 | 1.7 | 0.9 |
| | | Average | 2.2 | 5.1 | 4.1 | 0.6 | 7.6 | 7.0 | 3.8 |
| | | Diff MInt-HDX | N/A | -3.0 | -1.9 | 1.6 | -5.4 | -4.8 | -1.6 |
| | Rep 2 | Pose 1 | 1.7 | 2.2 | 2.6 | 0.6 | 10.0 | 1.7 | 9.7 |
| | | Pose 2 | 1.9 | 10.0 | 2.5 | 0.6 | 10.0 | 9.2 | 0.9 |
| | | Pose 3 | 0.9 | 2.7 | 2.8 | 0.6 | 2.7 | 1.9 | 9.2 |
| | | Average | 1.5 | 5.0 | 2.6 | 0.6 | 7.6 | 4.3 | 6.6 |
| | | Diff MInt-HDX | N/A | -3.5 | -1.2 | 0.9 | -6.1 | -2.8 | -5.1 |
| | Rep 3 | Pose 1 | 1.7 | 2.2 | 2.1 | 0.6 | 1.1 | 1.7 | 2.8 |
| | | Pose 2 | 3.1 | 9.9 | 2.6 | 0.6 | 10.0 | 9.2 | 2.7 |
| | | Pose 3 | 0.9 | 4.0 | 2.6 | 0.6 | 10.0 | 9.6 | 0.9 |
| | | Average | 1.9 | 5.4 | 2.4 | 0.6 | 7.1 | 6.8 | 2.1 |
| | | Diff MInt-HDX | N/A | -3.5 | -0.5 | 1.3 | -5.2 | -4.9 | -0.2 |
| | **Average Top Pose** | | **2.0** | **4.8** | **2.3** | **0.6** | **7.1** | **4.3** | **7.3** |
| | **Standard Deviation** | | **0.8** | **3.6** | **2.4** | **0.0** | **3.9** | **4.1** | **4.0** |
| | **Standard Deviation (Top-1)** | | **0.5** | **4.4** | **0.3** | **0.0** | **5.1** | **4.6** | **3.9** |
| | **Average Top 3 Poses** | | **1.8** | **5.2** | **3.0** | **0.6** | **7.4** | **6.0** | **4.2** |
| | **Average** | **Difference to MInt-HDX Top 3** | **-3.3** | **-1.2** | **1.3** | **-5.6** | **-4.2** | **-2.3** | |
| | | **Difference to MInt-HDX Top 1** | **-2.8** | **-0.3** | **1.4** | **-5.1** | **-2.3** | **-5.3** | |
| ERK2 | Rep 1 | Pose 1 | 3.1 | 10.4 | 5.7 | 3.5 | 3.1 | 3.2 | 10.5 |
| | | Pose 2 | 4.4 | 12.7 | 22.7 | 3.5 | 6.0 | 11.2 | 3.1 |
| | | Pose 3 | 4.1 | 9.8 | 3.1 | 3.4 | 5.8 | 11.3 | 4.7 |
| | | Average | 3.9 | 11.0 | 10.5 | 3.4 | 5.0 | 8.6 | 6.1 |
| | | Diff MInt-HDX | N/A | -7.1 | -6.6 | 0.4 | -1.1 | -4.7 | -2.2 |
| | Rep 2 | Pose 1 | 3.1 | 10.3 | 5.6 | 3.4 | 3.2 | 4.0 | 3.2 |
| | | Pose 2 | 4.3 | 9.1 | 23.1 | 3.4 | 6.0 | 3.4 | 11.0 |
| | | Pose 3 | 4.7 | 10.5 | 10.0 | 3.4 | 5.8 | 6.2 | 4.2 |
| | | Average | 4.0 | 10.0 | 12.9 | 3.4 | 5.0 | 4.5 | 6.1 |
| | | Diff MInt-HDX | N/A | -6.0 | -8.8 | 0.6 | -1.0 | -0.5 | -2.1 |
| | Rep 3 | Pose 1 | 3.1 | 12.5 | 33.0 | 3.7 | 3.2 | 3.2 | 11.0 |
| | | Pose 2 | 3.3 | 8.9 | 5.6 | 3.4 | 6.0 | 11.2 | 4.3 |
| | | Pose 3 | 4.8 | 12.7 | 10.2 | 3.3 | 5.8 | 3.1 | 10.9 |
| | | Average | 3.7 | 11.3 | 16.3 | 3.5 | 5.0 | 5.8 | 8.7 |
| | | Diff MInt-HDX | N/A | -7.6 | -12.5 | 0.2 | -1.3 | -2.1 | -5.0 |
| | **Average Top Pose** | | **3.1** | **11.1** | **14.8** | **3.6** | **3.1** | **3.5** | **8.2** |
| | **Standard Deviation** | | **0.7** | **1.5** | **10.5** | **0.1** | **1.4** | **3.8** | **3.7** |
| | **Standard Deviation (Top-1)** | | **0.0** | **1.2** | **15.8** | **0.1** | **0.0** | **0.5** | **4.4** |
| | **Average Top 3 Poses** | | **3.9** | **10.8** | **13.2** | **3.4** | **5.0** | **6.3** | **7.0** |
| | **Average** | **Difference to MInt-HDX Top 3** | **-6.9** | **-9.3** | **0.4** | **-1.1** | **-2.4** | **-3.1** | |
| | | **Difference to MInt-HDX Top 1** | **-7.9** | **-11.6** | **-0.4** | **0.0** | **-0.3** | **-5.1** | |
| FosA | Rep 1 | Pose 1 | 2.1 | 19.7 | 3.3 | 2.4 | 7.0 | 9.1 | 5.7 |
| | | Pose 2 | 3.1 | 20.3 | 12.2 | 2.3 | 6.3 | 2.1 | 6.3 |
| | | Pose 3 | 4.0 | 19.7 | 6.7 | 2.1 | 7.1 | 2.9 | 7.2 |
| | | Average | 3.1 | 19.9 | 7.4 | 2.3 | 6.8 | 4.7 | 6.4 |
| | | Diff MInt-HDX | N/A | -16.8 | -4.3 | 0.8 | -3.7 | -1.6 | -3.3 |
| | Rep 2 | Pose 1 | 2.1 | 19.9 | 19.7 | 2.2 | 7.0 | 9.1 | 5.5 |
| | | Pose 2 | 3.8 | 20.4 | 20.0 | 2.1 | 6.4 | 2.1 | 6.3 |
| | | Pose 3 | 3.9 | 19.7 | 20.1 | 2.3 | 7.1 | 10.2 | 7.2 |
| | | Average | 3.3 | 20.0 | 19.9 | 2.2 | 6.9 | 7.1 | 6.3 |
| | | Diff MInt-HDX | N/A | -16.7 | -16.6 | 1.1 | -3.6 | -3.8 | -3.1 |
| | Rep 3 | Pose 1 | 2.1 | 19.7 | 19.7 | 1.9 | 7.0 | 9.1 | 5.7 |
| | | Pose 2 | 3.6 | 19.7 | 21.2 | 2.2 | 6.5 | 2.2 | 6.3 |
| | | Pose 3 | 4.1 | 20.4 | 21.0 | 2.2 | 6.7 | 3.7 | 7.2 |
| | | Average | 3.3 | 19.9 | 20.6 | 2.1 | 6.7 | 5.0 | 6.4 |
| | | Diff MInt-HDX | N/A | -16.7 | -17.4 | 1.1 | -3.5 | -1.7 | -3.1 |
| | **Average Top Pose** | | **2.1** | **19.8** | **14.2** | **2.2** | **7.0** | **9.1** | **5.6** |
| | **Standard Deviation** | | **0.9** | **0.3** | **6.8** | **0.1** | **0.3** | **3.6** | **0.7** |
| | **Standard Deviation (Top-1)** | | **0.0** | **0.1** | **9.5** | **0.2** | **0.0** | **0.0** | **0.1** |
| | **Average Top 3 Poses** | | **3.2** | **19.9** | **16.0** | **2.2** | **6.8** | **5.6** | **6.4** |
| | **Average** | **Difference to MInt-HDX Top 3** | **-16.7** | **-12.8** | **1.0** | **-3.6** | **-2.4** | **-3.2** | |
| | | **Difference to MInt-HDX Top 1** | **-17.7** | **-12.1** | **0.0** | **-4.9** | **-6.9** | **-3.5** | |
| | **Overall Average** | | **3.0** | **12.0** | **10.8** | **2.1** | **6.4** | **6.0** | **5.9** |
| | **Standard Deviation** | | **0.8** | **1.8** | **6.6** | **0.1** | **1.9** | **3.8** | **2.8** |
| | **Top 3 Pose Diff Avg** | | **N/A** | **-9.0** | **-7.8** | **0.9** | **-3.4** | **-3.0** | **-2.9** |
| | **Top Pose Overall Avg** | | **2.4** | **11.9** | **10.4** | **2.1** | **5.8** | **5.6** | **7.1** |
| | **Standard Deviation (Top-1)** | | **0.6** | **2.2** | **9.4** | **0.3** | **2.5** | **2.2** | **2.9** |
| | **Top Pose Diff Avg** | | **N/A** | **-9.5** | **-8.0** | **0.3** | **-3.3** | **-3.2** | **-4.6** |

**Supplemental Table S10: Validation set L-RMSD across replicas and methods.** The L-RMSD for each protein in the validation set, across each replicate is shown for the Top-3 poses as determined by each respective tool and for the reduced data set to probe the effect of HDX-MS data quality factors on MInt-HDX. For each protein, the average Top-1 and Top-3 poses are calculated and shown, as well as the overall average across all proteins.

| Peptide Segment Length | | | | | | | | |
|---|---|---|---|---|---|---|---|---|
| System | Total Coverage | Data Split >9/<10 | Average Peptide Length >9 | Average Peptide Length <10 | >9 Coverage | <10 Coverage | Redundancy >9 | Redundancy <10 |
| VDR | 93% | 25% / 75% | 13.8 | 6 | 53% | 81% | 3.07 | 2.73 |
| ERK2 | 90% | 55% / 45% | 14.3 | 7.4 | 77% | 50% | 2.51 | 1.71 |
| FosA | 99% | 40% / 60% | 12.6 | 6.3 | 78% | 80% | 3.06 | 2.55 |
| **Average** | 94% | 40% / 60% | 13.6 | 6.6 | 69% | 70% | 2.9 | 2.3 |
| **Peptide Segment Redundancy** | | | | | | | | |
| System | Total Redundancy | Total Coverage | Total Average Peptide Length | % Total Data | Redundancy | % Coverage | Average Peptide Length | |
| VDR | 3.99 | 93% | 7.9 | 29.7% | 1.11 | 91% | 6.9 | |
| ERK2 | 3.07 | 90% | 11.2 | 40.8% | 1.17 | 81% | 10.7 | |
| FosA | 4.45 | 99% | 8.8 | 26.0% | 1.11 | 98% | 8.3 | |
| **Average** | 3.84 | 94% | 9.3 | 32.20% | 1.13 | 90% | 8.6 | |

**Supplemental Table S11: HDX-MS peptide segment length, redundancy, and coverage.** For the peptide segment length and peptide redundancy tests, the coverage, data split, average length, and redundancy was calculated for each validation set protein and each artificial data split.

| System | Replicate | | MInt-HDX L-RMSD |
|---|---|---|---|
| DrDXPS AF3 Conf 0.5 | Rep 1 | Pose 1 | 1.7 |
| | | Pose 2 | 2.0 |
| | | Pose 3 | 1.5 |
| | | Average | 1.7 |
| | Rep 2 | Pose 1 | 1.7 |
| | | Pose 2 | 2.0 |
| | | Pose 3 | 1.8 |
| | | Average | 1.9 |
| | Rep 3 | Pose 1 | 1.8 |
| | | Pose 2 | 2.0 |
| | | Pose 3 | 1.9 |
| | | Average | 1.9 |
| | **Average Top Pose** | | **1.7** |
| | **Standard Deviation** | | **0.2** |
| | **Standard Deviation (Top-1)** | | **0.01** |
| | **Average Top 3 Poses** | | **1.8** |
| DrDXPS AF3 Conf 0.7 | Rep 1 | Pose 1 | 19.5 |
| | | Pose 2 | 20.4 |
| | | Pose 3 | 20.4 |
| | | Average | 20.1 |
| | Rep 2 | Pose 1 | 19.4 |
| | | Pose 2 | 20.5 |
| | | Pose 3 | 17.7 |
| | | Average | 19.2 |
| | Rep 3 | Pose 1 | 19.4 |
| | | Pose 2 | 20.3 |
| | | Pose 3 | 18.9 |
| | | Average | 19.5 |
| | **Average Top Pose** | | **19.4** |
| | **Standard Deviation** | | **0.9** |
| | **Standard Deviation (Top-1)** | | **0.06** |
| | **Average Top 3 Poses** | | **19.6** |

**Supplemental Table S12: L-RMSD of replicas of MAP docking to *Dr*DXPS at confidence threshold 0.5 and 0.7.** Shown is the L-RMSD of each individual replicate's Top-3 poses for *Dr*DXPS with a confidence threshold value of 0.5 and then 0.7.

| System | System Docked To | | L-RMSD to Chicken DFG Out (2OIQ, Chain A) | L-RMSD to Human Analog DFG In (1Y57) | Score |
|---|---|---|---|---|---|
| CSRC | Chicken DFG Out Rep 1 (PDB: 2OIQ, Chain A) | Pose 1 | 1.6 | 11.5 | 5.7 |
| | | Pose 2 | 2.1 | 10.8 | 5.0 |
| | | Pose 3 | 8.2 | 5.8 | 5.0 |
| | | Average | 4.0 | 9.4 | 5.2 |
| | Chicken DFG Out Rep 2(PDB: 2OIQ, Chain A) | Pose 1 | 2.8 | 10.2 | 5.3 |
| | | Pose 2 | 3.1 | 10.6 | 5.3 |
| | | Pose 3 | 1.6 | 11.6 | 5.0 |
| | | Average | 2.5 | 10.8 | 5.2 |
| | Chicken DFG Out Rep 3 (PDB: 2OIQ, Chain A) | Pose 1 | 1.6 | 11.6 | 5.7 |
| | | Pose 2 | 2.1 | 10.9 | 5.0 |
| | | Pose 3 | 11.0 | 7.6 | 5.0 |
| | | Average | 4.9 | 10.0 | 5.2 |
| | Chicken DFG In Rep 1 (PDB: 2OIQ, Chain B) | Pose 1 | 10.9 | 2.7 | 4.0 |
| | | Pose 2 | 14.8 | 9.1 | 4.0 |
| | | Pose 3 | 10.3 | 4.1 | 4.0 |
| | | Average | 12.0 | 5.3 | 4.0 |
| | Chicken DFG In Rep 2 (PDB: 2OIQ, Chain B) | Pose 1 | 13.2 | 10.9 | 4.3 |
| | | Pose 2 | 10.9 | 2.6 | 4.3 |
| | | Pose 3 | 14.3 | 8.7 | 4.0 |
| | | Average | 12.8 | 7.4 | 4.2 |
| | Chicken DFG In Rep 3 (PDB: 2OIQ, Chain B) | Pose 1 | 10.9 | 2.7 | 3.7 |
| | | Pose 2 | 14.7 | 9.0 | 3.7 |
| | | Pose 3 | 10.0 | 5.8 | 3.7 |
| | | Average | 11.9 | 5.8 | 3.7 |
| | Human DFG In Bound Analog Rep 1 (PDB: 1Y57) | Pose 1 | 12.8 | 6.2 | 4.0 |
| | | Pose 2 | 11.8 | 7.3 | 4.0 |
| | | Pose 3 | 12.5 | 7.1 | 4.0 |
| | | Average | 12.4 | 6.9 | 4.0 |
| | Human DFG In Bound Analog Rep 2 (PDB: 1Y57) | Pose 1 | 11.8 | 7.3 | 4.0 |
| | | Pose 2 | 12.8 | 6.2 | 4.0 |
| | | Pose 3 | 12.4 | 7.2 | 4.0 |
| | | Average | 12.3 | 6.9 | 4.0 |
| | Human DFG In Bound Analog Rep 3 (PDB: 1Y57) | Pose 1 | 12.9 | 6.2 | 4.0 |
| | | Pose 2 | 11.8 | 7.3 | 4.0 |
| | | Pose 3 | 12.8 | 7.3 | 4.0 |
| | | Average | 12.5 | 6.9 | 4.0 |

| System | System Docked To | Average Top 1 MInt-HDX Score | Average Top 3 MInt-HDX Score | Average Top 1 L-RMSD | Average Top 3 L-RMSD |
|---|---|---|---|---|---|
| CSRC | Chicken DFG Out (PDB: 2OIQ, Chain A) | 5.6 | 5.2 | 2.0 | 3.8 |
| | Chicken DFG In (PDB: 2OIQ, Chain B) | 4.0 | 4.0 | 5.4 | 6.2 |
| | Human DFG In Bound Analog (PDB: 1Y57) | 4.0 | 4.0 | 6.6 | 6.9 |

**RMSD (Angstroms) of the DFG Region**

| System | Chicken c-src (PDB: 2OIQ, Chain A)- DFG-out | Chicken c-src (PDB: 2OIQ, Chain B) | Human c-src (PDB: 1Y57) DFG-In |
|---|---|---|---|
| Chicken c-src (PDB: 2OIQ, Chain A)- DFG-out | 0.0 | | |
| Chicken c-src (PDB: 2OIQ, Chain B) | 6.7 | 0.0 | |
| Human c-src (PDB: 1Y57) DFG-In | 6.5 | 1.2 | 0.0 |

**Supplemental Table S13: C-src kinase replica MInt-HDX docking, L-RMSD and scores, and crystal structure RMSD's of the DFG region and ligand. Top table-** For each DFG conformation of c-src kinase (DFG-in 1Y57 or 2OIQ Chain B vs DFG-out 2OIQ Chain A), the L-RMSD of docked Imatinib is shown to the ligand conformation in 2OIQ Chain A (Imatinib) or in 1Y57 (MPZ). The last column displays the score as determined by MInt-HDX. **Middle table-** The average Top-1 and Top-3 MInt-HDX scores and L-RMSDs are shown. **Bottom Table-** The pairwise all-atom RMSD of the DFG regions in the input structures are shown after structural alignment using backbone atoms of the entire protein.

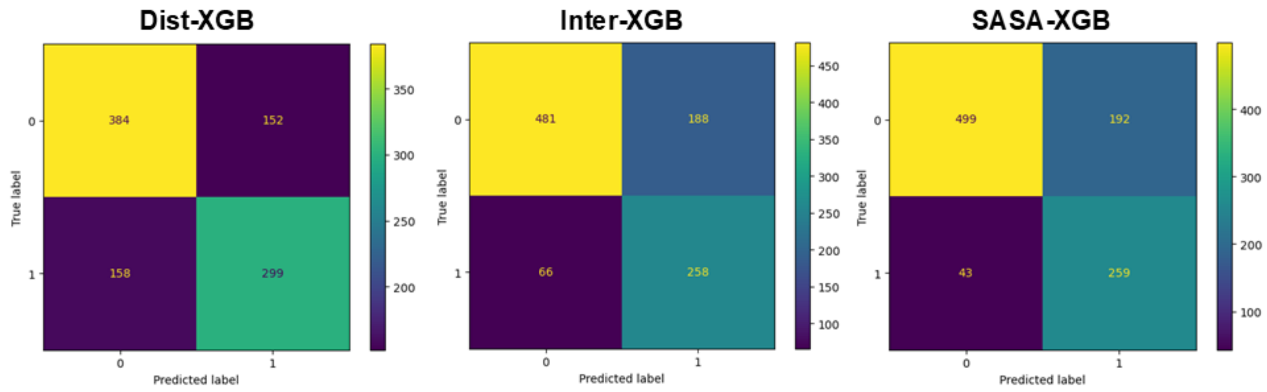


**Supplemental Figure 1: XGBoost models performance on the validation set data.** Shown above is the confusion matrix for the three different XGBoost models trained, going from distance, to interactions, and then SASA respectively. On the Y-axis are the true labels as derived from the bound crystal structure of the protein, and on the X-axis are the predicted label as determined by the XGBoost model.

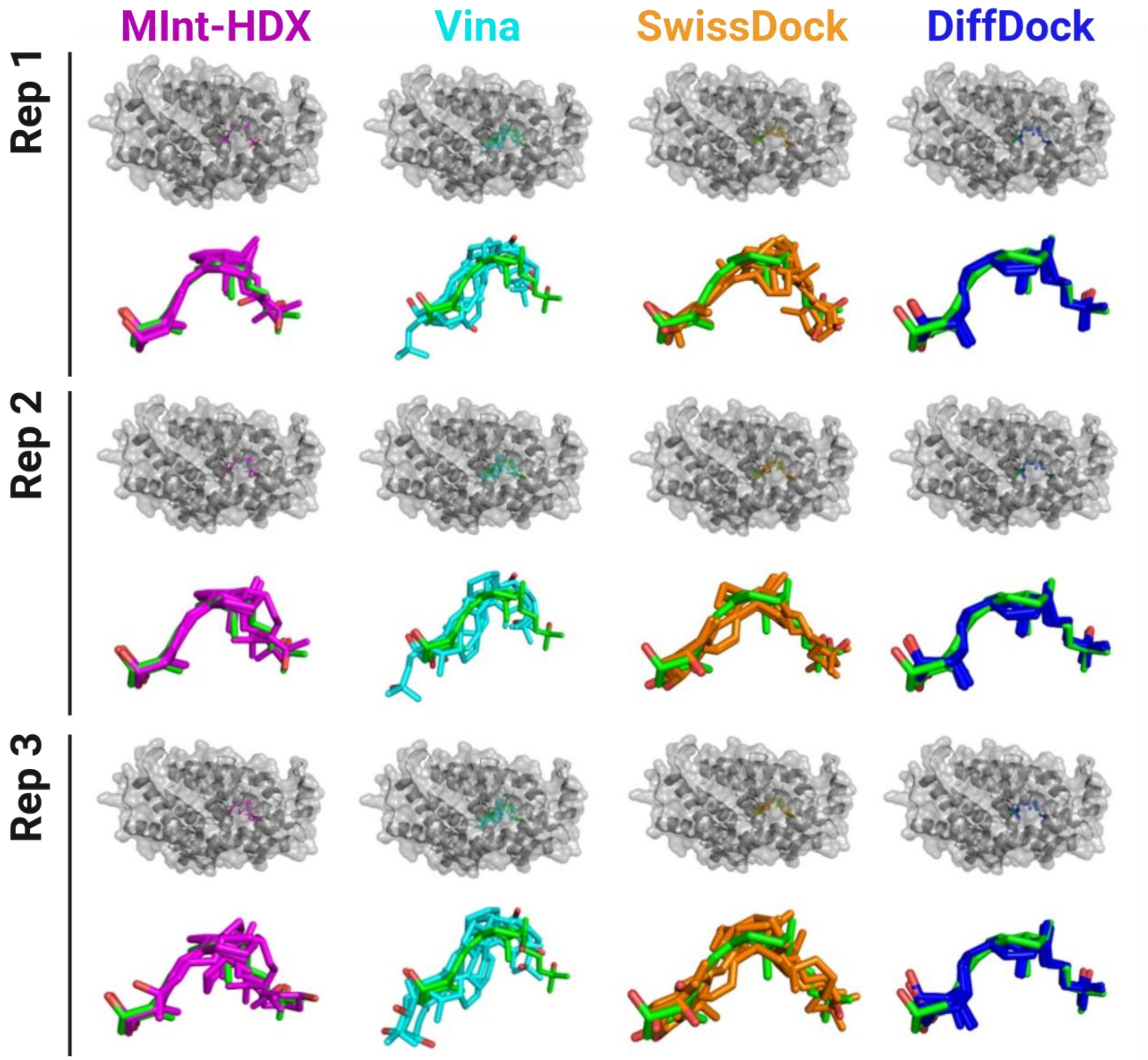


**Supplemental Figure 2: Three docking replicates of each docking tool to VDR.** The docking poses from MInt-HDX (Magenta), AutoDock Vina (Cyan), SwissDock (Orange), and DiffDock (Blue) were aligned to the ligand from the bound crystal structure (Green). Alignment is performed based on protein backbone atoms. Shown are the Top-3 docking poses of the three replicates for each tool when docking to the protein VDR (PDB: 1DB1). The top shows the zoomed out, full protein view with the ligands in sticks and protein in surface representation. The bottom images illustrate the ligand conformation zoomed in, without the protein.

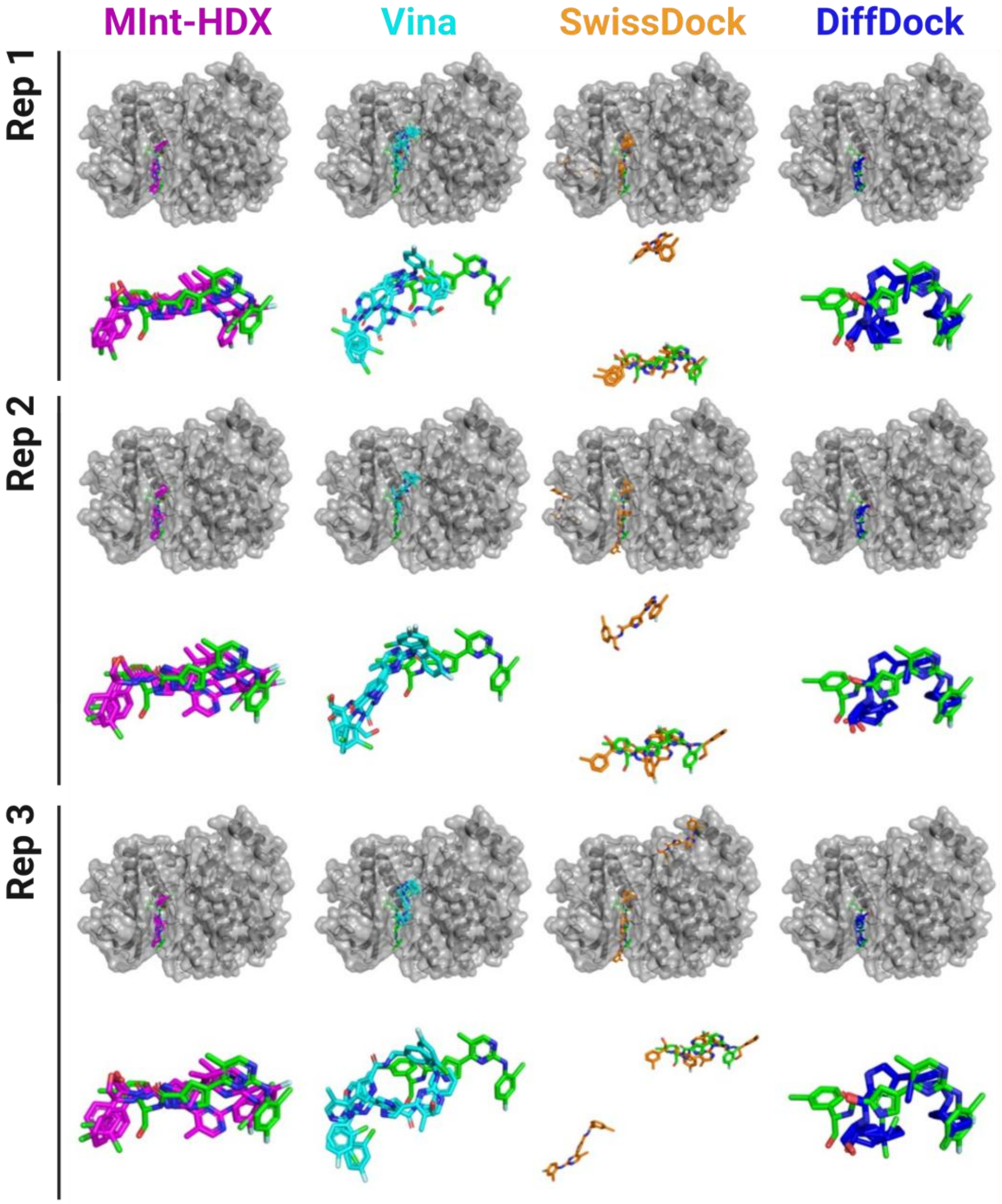


**Supplemental Figure 3: Three docking replicates of each tool to ERK2** The docking poses from MInt-HDX (Magenta), AutoDock Vina (Cyan), SwissDock (Orange), and DiffDock (Blue) were aligned to the ligand from the bound crystal structure (Green). Alignment is performed based on protein backbone atoms. Shown are the Top-3 docking poses of the three replicates for each tool when docking to the protein ERK2 (PDB: 6OPK). The top shows the zoomed out, full protein view with the ligands in sticks and protein in surface representation. The bottom images illustrate the ligand conformation zoomed in, without the protein.

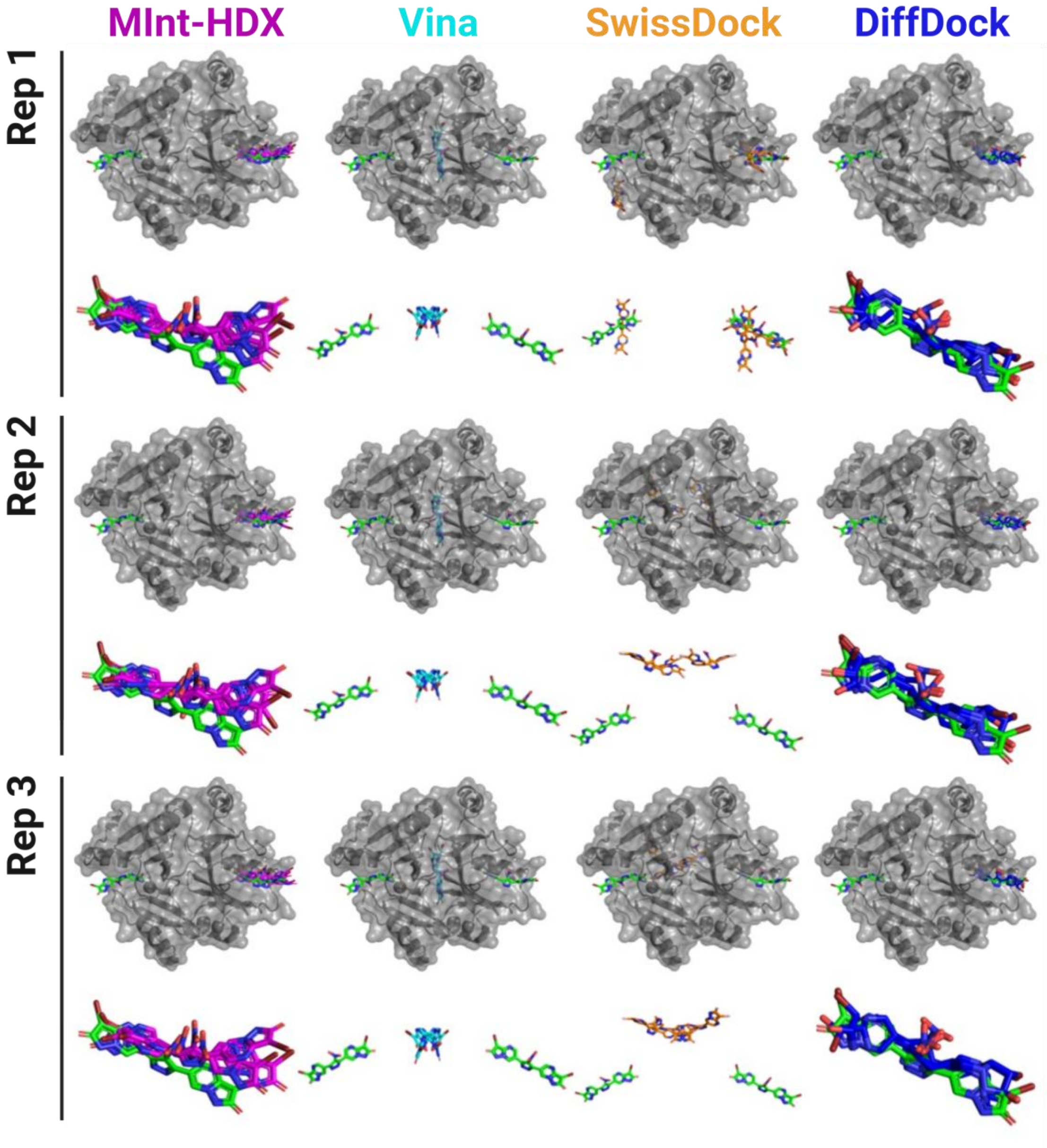


**Supplemental Figure 4: Three docking replicates of each tool to FOSA** The docking poses from MInt-HDX (Magenta), AutoDock Vina (Cyan), SwissDock (Orange), and DiffDock (Blue) were aligned to the ligand from the bound crystal structure (Green). Alignment is performed based on protein backbone atoms. Shown are the Top-3 docking poses of the three replicates for each tool when docking to the protein FOSA (PDB: 5WEW). The top shows the zoomed out, full protein view with the ligands in sticks and protein in surface representation. The bottom images illustrate the ligand conformation zoomed in, without the protein. For MInt-HDX in Magenta and DiffDock in Blue, the zoomed in view of the ligands, due to the predictions only being to one part of the homodimer, has the second ligand omitted.

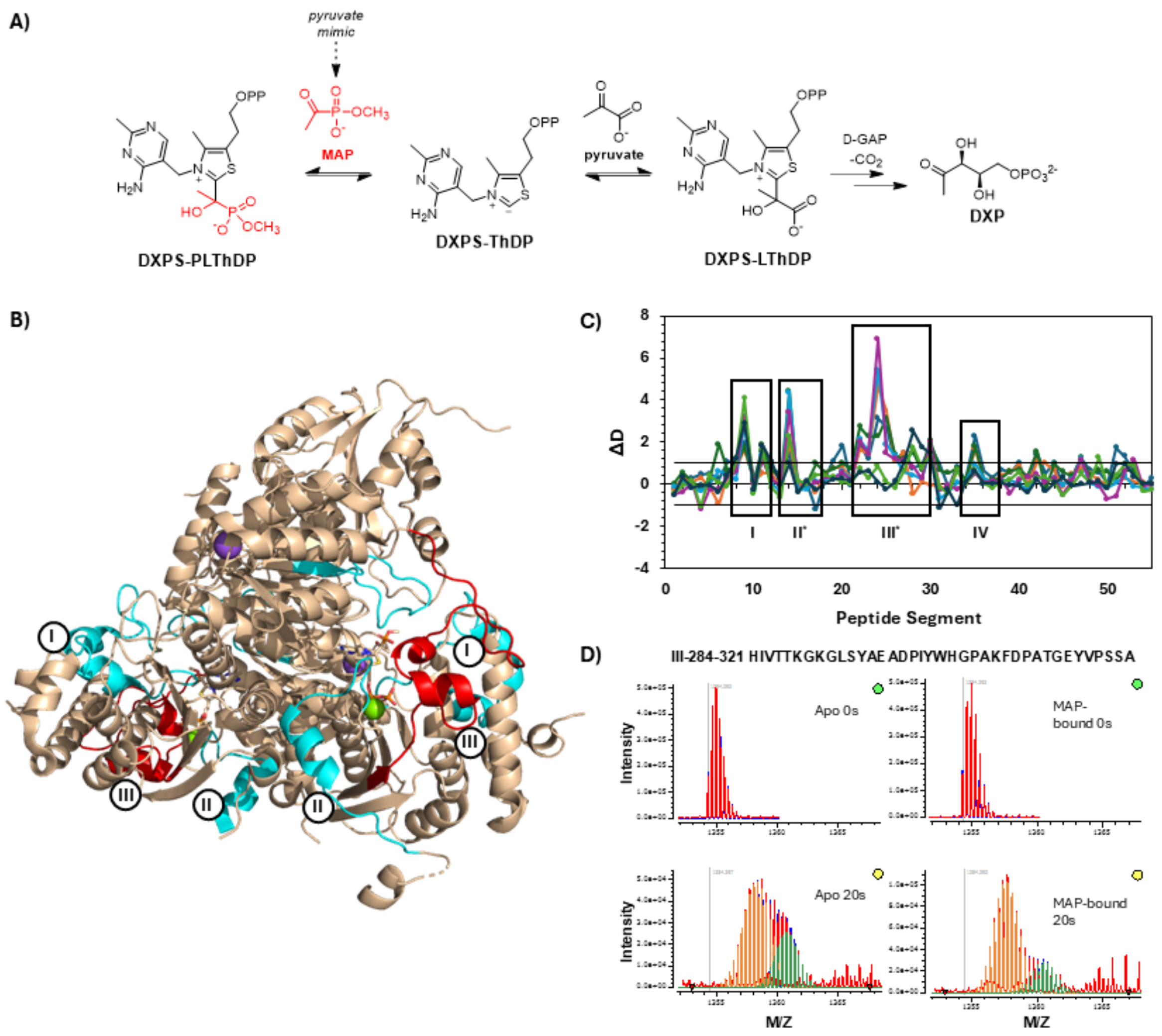


**Supplemental Figure 5: HDX-MS data for *Dr*DXPS.** HDX-MS Data for the *Dr*DXPS enzyme with and without MAP was acquired. Four main regions were identified due to their differences in deuteration between the bound and unbound states. **A)** Scheme of the DXPS catalyzed formation of the LThPD ligand with pyruvate as a substrate and the inhibitory PLThDP ligand with MAP as pyruvate mimicking substrate. MAP atoms are shown in red. **B)** Represents the four regions mapped to the protein's crystal structure (PDB: 6OUV). The three regions colored in cyan correspond to an increased level of protection from exchange as observed by the HDX-MS data. The third region, colored in red, is indicative of the EX1 Kinetic region. **C)** Difference plot showing the difference in hydrogen-deuterium exchange between the bound and unbound states. Regions mapped in Panel A have been annotated accordingly. **D)** Mass spectra for peptide 284-321 showing bimodal isotopic envelops consistent with EX1 Kinetics in both the bound and unbound form.

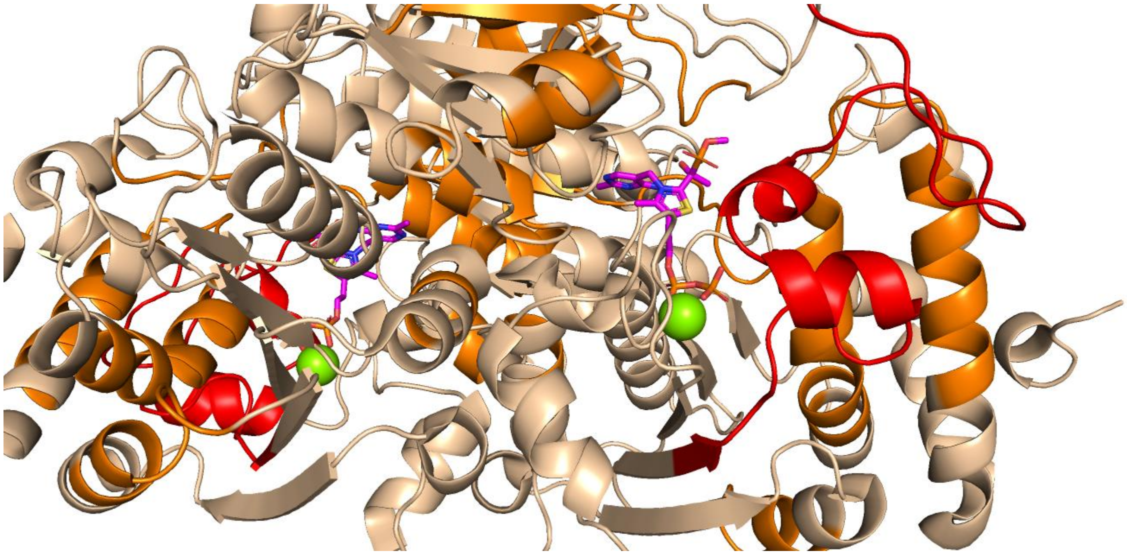

**Supplemental Figure 6: Areas of missing HDX-MS coverage mapped to the crystal structure of *Dr*DXPS.** Shown in orange are the areas of missing HDX-MS data coverage, mapped to the bound crystal structure of the protein (PDB: 6OUV). The area shown in red is the region of the protein displaying EX1 kinetics (residues 284-321).

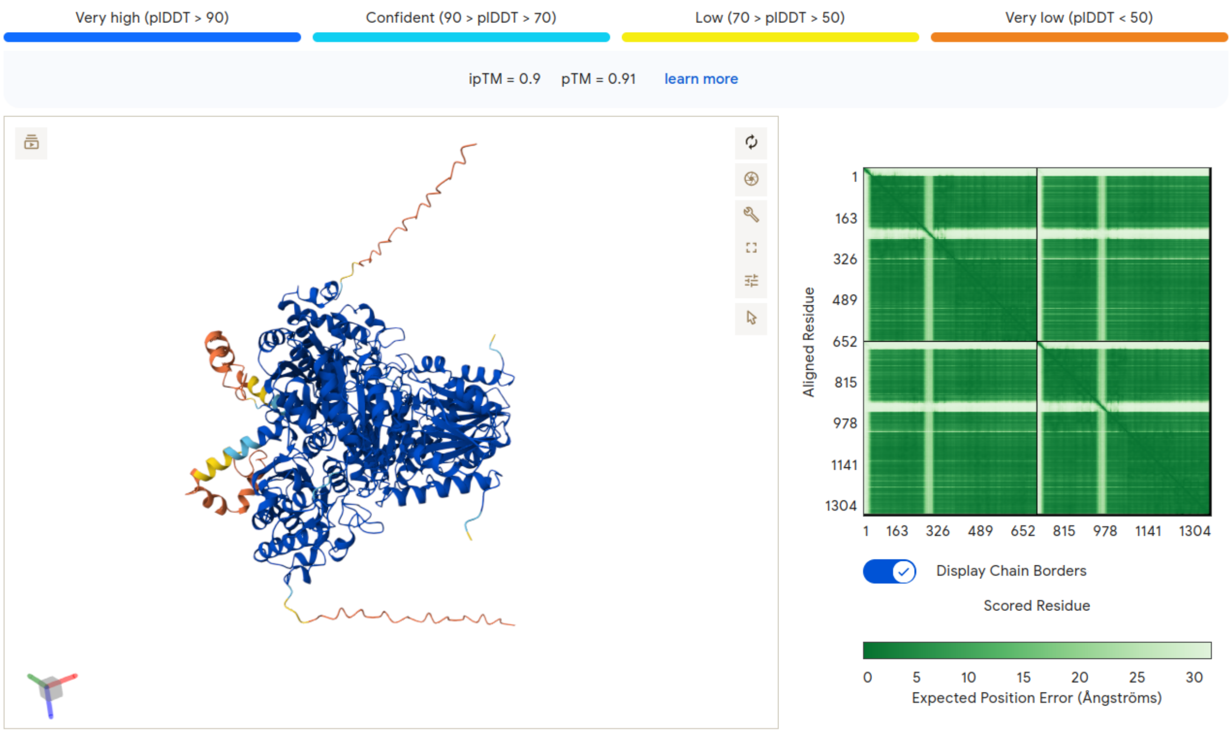


**Supplemental Figure 7: AlphaFold III model confidence map for the apo form of *Dr*DXPS.** The AlphaFold III modeled structure has high confidence for the majority of the protein (blue and cyan). Areas of missing electron density in the bound crystal structure can be seen to have lower confidence (yellow and orange). The binding site is located within the region of high confidence.
